# Nanopore sequencing of intact aminoacylated tRNAs

**DOI:** 10.1101/2024.11.18.623114

**Authors:** Laura K. White, Aleksandar Radakovic, Marcin P. Sajek, Kezia Dobson, Kent A. Riemondy, Samantha del Pozo, Jack W. Szostak, Jay R. Hesselberth

## Abstract

Transfer RNAs (tRNA) are decorated during biogenesis with a variety of modifications that modulate their stability, aminoacylation, and decoding potential during translation. The complex landscape of tRNA modification presents significant analysis challenges and to date no single approach enables the simultaneous measurement of important but disparate chemical properties of individual, mature tRNA molecules. We developed a new, integrated approach to analyze the sequence, modification, and aminoacylation state of tRNA molecules in a high throughput nanopore sequencing experiment, leveraging a chemical ligation that embeds the charged amino acid in an adapted tRNA molecule. During nanopore sequencing, the embedded amino acid generates unique distortions in ionic current and translocation speed, enabling application of machine learning approaches to classify charging status and amino acid identity. Specific applications of the method indicate it will be broadly useful for examining relationships and dependencies between tRNA sequence, modification, and aminoacylation.

## INTRODUCTION

Transfer RNAs (tRNAs) are the fundamental adaptors of translation, linking mRNA decoding and polypeptide synthesis (*1*). Biogenesis of tRNAs is complex: following transcription, precursor tRNA molecules undergo multiple trimming and modification events, yielding small (∼76 nt) and densely modified (∼13 modifications) RNA molecules that are then competent for aminoacylation (reviewed in (*2*)). tRNA modifications can profoundly impact tRNA function through their impacts on stability (*3*), aminoacylation efficiency (*4*, *5*), and expanded decoding capacity (*6*). tRNA charging levels vary in response to environmental conditions (*7*, *8*) and drive various physiological responses to stress (*9*), starvation (*10*, *11*), and proliferation (*12*, *13*). In addition, aminoacylation of tRNAs with non-cognate amino acids (“misaminoacylation”) has been observed at high rates in aminoacyl synthetases with proofreading defects and during oxidative stress (*14–18*).

High throughput analysis of tRNAs has been hampered by a lack of incisive methods to comprehensively and directly measure tRNA abundance, modification, and aminoacylation within the same experiment. Short-read DNA sequencing approaches have been developed to study tRNA charging, but begin by selective oxidation and β-elimination of non-aminoacylated tRNAs, marking this population through removal of the 3′-terminal nucleotide, followed by deacylation, adapter ligation, and reverse transcription (*19–21*). However, the large number of modifications installed on tRNAs make them poor substrates for reverse transcriptase (RT) enzymes. In addition, most tRNA modifications do not produce detectable RT signatures (*22*) or require chemical derivatization to do so (*23*). As such, current sequencing approaches for studying aminoacylated tRNAs rely on indirect information to analyze key properties of these molecules, using chemical and enzymatic treatments to infer the positions of body modifications and presence of charged amino acids. Hence, lower throughput and more cumbersome techniques capable of physically separating uncharged tRNA and aminoacylated tRNA (aa-tRNA) (*24*, *25*) remain the gold standard for quantification of tRNA aminoacylation.

A promising new direction is the application of nanopores to tRNA sequencing. Successful discrimination of several tRNAs by solid-state (*26*) and biological (*27*, *28*) nanopores has been followed by methods to directly sequence tRNAs on a commercially available nanopore platform (*29*), enabling direct analysis of the RNA molecule (*30*). Further refinement in adapter design and computational analysis have improved tRNA nanopore sequencing to the point that it is now possible to directly assess tRNA abundance and modification status with single molecule precision (*31*, *32*).

Here, we develop a nanopore sequencing approach (“aa-tRNA-seq”) that directly captures information on tRNA sequence, modifications, and aminoacylation in a single read. Our method enables selective capture of tRNAs based on their aminoacylation status, selectively embedding the amino acid of aa-tRNAs within the adapter-ligated tRNA molecule and separately capturing non-aminoacylated tRNA, facilitating comparative analyses of tRNA charging. We have characterized nanopore signals produced by 20 proteinogenic amino acids using synthetic tRNA, and leveraged these signals to train a convolutional neural network (CNN) to discriminate aminoacylated tRNAs from their uncharged counterparts, and extended this approach for pairwise amino acid classification. We applied the method to study changes in budding yeast tRNA populations during nutrient limitation and in conditions of tRNA hypomodification, confirming known and identifying unexpected changes in tRNA aminoacylation and abundance during these stress conditions.

## RESULTS

### A chemical ligation approach enables selective capture of intact aminoacylated tRNAs

While investigating prebiotic roles of aminoacyl-RNAs, we developed a splinted ligation reaction that generates amino acid-bridged chimeric RNA molecules using a 5′-phosphorimidazole activated oligoribonucleotide (*33*, *34*). We realized that aminoacyl-tRNAs were analogous to these substrates, so we tested and found that a synthetic tRNA-Gly-GCC aminoacylated with glycine or lysine using the Flexizyme (*35*) underwent chemical ligation with moderate kinetics, while the non-aminoacylated tRNA yielded no detectable product (**Fig. S1A**). To accelerate the reaction between the ɑ-amino group of the aminoacyl-tRNA and the activated adapter, we included 1-(2-Hydroxyethyl)imidazole (HEI) as an organocatalyst (*36*) (**Fig. 1A, S1B**), and subjected *S. cerevisiae* tRNA to a 30 minute chemical ligation at pH 5.5. Under these conditions, we achieved efficient ligation of aminoacylated budding yeast tRNA as measured by acidic northern blot, with <0.1% background ligation to a chemically deacylated tRNA control for tRNA-Gly-GCC (**Fig. 1B**). To examine the efficiency of this reaction on biological tRNA substrates, we stripped and reprobed the same membrane from **Fig. 1B** using oligonucleotide probes complementary to isodecoders from each of the 20 tRNA isoacceptor families in yeast. Of the 16 tRNA isodecoders that were separated sufficiently to enable densitometric quantification, the percent of aminoacylated species shifted upon chemical ligation ranged from 62-100%, with quantitative ligation of arginyl, asparaginyl, cysteinyl, glutaminyl, glycyl, and lysyl tRNA species (**Fig. S1D**). Using a fluorescently labeled tRNA analog, we confirmed this HEI-catalyzed reaction yields nearly quantitative ligation *in vitro* to substrates that were Flexizyme-charged with 13 different amino acids (**Fig. S1B,C**).

**Figure 1.**
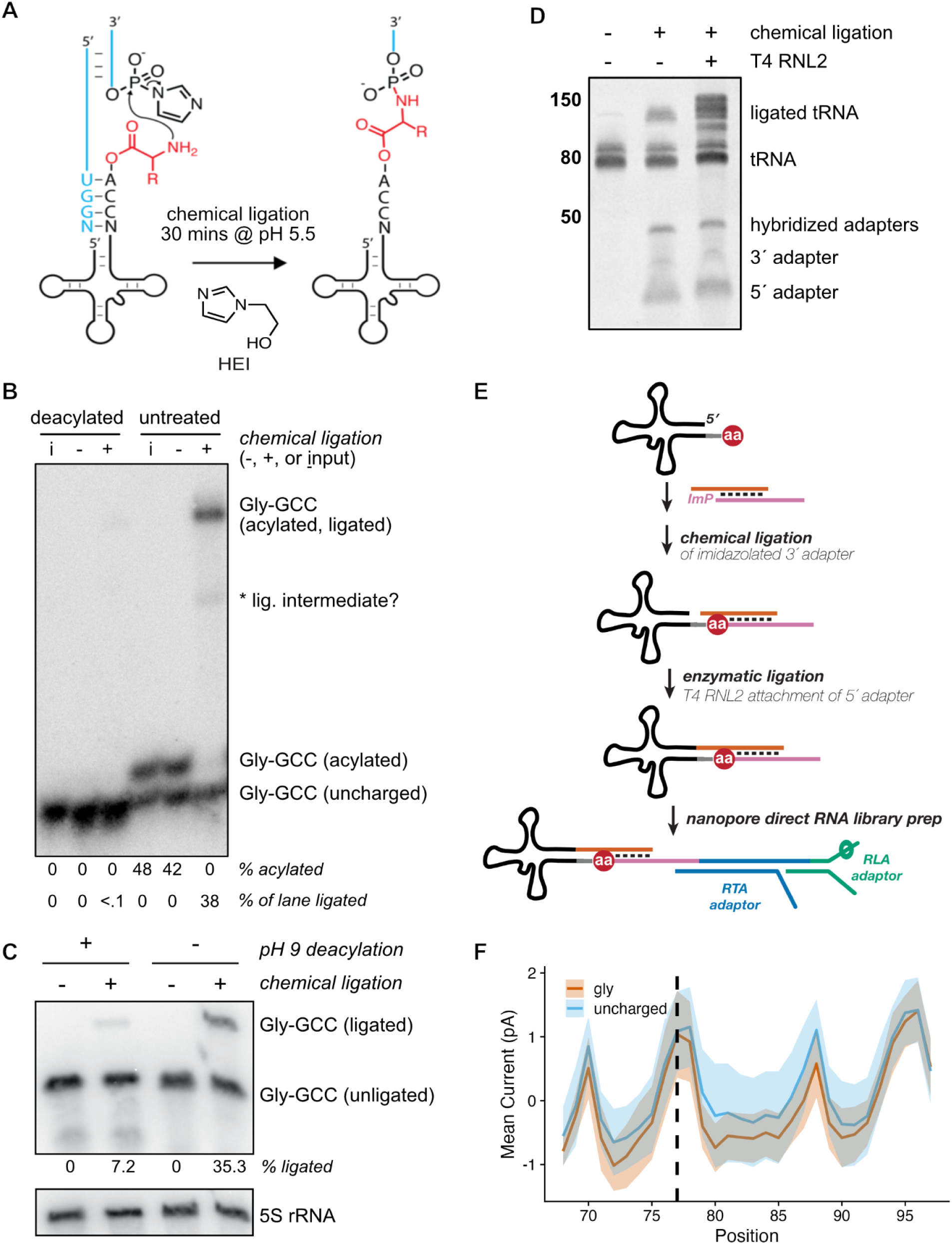
A chemical ligation strategy for capture of aminoacylated tRNAs. (**A**) Schematic of splinted chemical ligation of 5′-phosphorimidazolide activated adapter (blue) to aminoacyl tRNA (black, with amino acid in red) in the presence of the catalyst 1-(2-Hydroxyethyl)imidazole (HEI). (**B**) Acidic charging northern of deacylated or untreated wild-type (BY4741) *S. cerevisiae* tRNA after chemical reaction in the presence (+) or absence (-) of activated 3′ adapter, compared to tRNA input only (I). Blot has been probed for budding yeast tRNA Gly-GCC; asterisk indicates a presumed ligation intermediate. (**C**) “Chemical-charging northern blot” resolving charged and uncharged tRNA-Gly-GCC via chemical ligation and analysis on a mini (8×10 cm) TBE / 7M urea polyacrylamide gel followed by membrane transfer and hybridization with a Gly-GCC probe. (**D**) Small RNAs (17-200 nt, 200 ng) from budding yeast were chemically ligated to activated 3′ RNA adapter in a splinted ligation as previous panels, followed by an optional enzymatic ligation with T4 RNA ligase 2 (RNL2) to attach a second, splinted RNA adapter to the 5′ end of the tRNA. Products were run on a 10% denaturing PAGE gel and stained with SYBR Gold. (**E**) Schematic illustrating strategy for chemical ligation and sequencing of charged and uncharged biological tRNAs via nanopore direct RNA sequencing. (**F**) Normalized mean current in picoamps for synthetic tRNA charged with glycine vs an uncharged control. The region visualized includes the 3′ terminus of the tRNA (6 nt, positions 68-73), the CCA tail (positions 74-76), the aminoacylated position (dashed line), and the entirety of the 3′ adapter sequence. For each colored trace, the solid line is the mean signal, and the shading spans the standard deviation.

### A chemical-northern blot simplifies analysis of aminoacylated tRNA

The analysis of tRNA aminoacylation by acidic northern blotting requires careful handling to preserve the labile ester linkage, using a large format, low pH polyacrylamide gel run in cold, acidic buffer conditions to provide adequate resolution of charged and uncharged tRNA, which is achieved by >12 hours of electrophoresis depending on the isodecoder of interest (*24*). Chemical ligation of aminoacyl-tRNA stabilizes the ester (*33*, *37*, *38*) and significantly increases its size, enabling robust separation from non-acylated tRNA by acidic northern blot (**Fig. 1B**). These features motivated a simpler approach to separate charged and uncharged tRNA using standard denaturing polyacrylamide gel electrophoresis. In this “chemical-charging northern” (**Fig. 1C**) budding yeast tRNA was chemically ligated as in **Fig. 1B**, followed by a ∼30 minute electrophoretic separation on a standard TBE-urea mini-gel, membrane transfer, and probe hybridization for the same glycyl isodecoder. We found that 35% of Gly-GCC tRNA displays a gel shift after chemical ligation (**Fig. 1C**, lane 4, consistent with the value from **Fig. 1B**), with 7% background ligation (due to incomplete deacylation) to the deacylated sample (lane 2). In the absence of the phosphorimidazole-activated 3′ oligoribonucleotide (lanes 1 & 3), no gel shift for the Gly-GCC tRNA is apparent.

### Aminoacylated tRNAs can be analyzed by nanopore sequencing

We next assessed whether synthetic tRNA ligated to an activated 3′ oligoribonucleotide via a bridging amino acid was amenable to nanopore sequencing. We confirmed that adapters for direct tRNA nanopore sequencing (*32*) can be attached to budding yeast tRNA via sequential chemical and enzymatic ligations of the 3′ and 5′ sequences, respectively (**Fig. 1D**), suggesting a clear strategy for the preparation of nanopore sequencing libraries containing aminoacylated tRNAs (**Fig. 1E**). We synthesized glycyl-tRNA charged with the Flexizyme (*35*), chemically ligated this to the same phosphorimidazolated 3′ adapter, gel purified the ligation product, and enzymatically ligated the 5′ adapter using T4 RNA ligase 2 (RNL2). We then prepared a nanopore direct RNA sequencing library from the individual ligated glycyl-tRNA using the RNA004 chemistry from Oxford Nanopore Technologies (ONT). **Fig. 1F** shows the difference in ionic current between a synthetic tRNA and the same sequence charged with glycine, with lower current for tRNA-Gly spanning multiple nucleotides as the aminoacylated molecule is pulled into the nanopore from its 3′ end during motor-catalyzed translocation.

We next synthesized tRNA charged with each of the 20 standard proteinogenic amino acids using the Flexizyme and subjected them to the same library preparation. Quality control assessment revealed no major issues for 18 of 20 of these libraries; however, 41.0% of asparaginyl tRNA reads and 27.6% of cysteinyl tRNA reads terminated aberrantly, consistent with these tRNAs becoming stuck in nanopores and subsequently ejected (“unblocked”) at higher rates than those charged with other, bulkier amino acids or the uncharged tRNA control (**Fig. S2A**). As the Asn-tRNA and Cys-tRNA libraries also had lower sequencing yields, we sought to resolve this issue and tested multiple strategies, illustrated in **Fig. S2B**. For tRNA-Cys, either (i) alkylation with chloroacetamide after chemical ligation or (ii) omitting the enzymatic ligation to the 5′ adapter resolved the pore blocking (**Fig. S2C**), suggesting that the thiol of Cys and amide of Asn might interact with the 5′ RNA adapter, leading to the molecule becoming stuck in the nanopore during sequencing. Indeed, changing the 5′ adapter from an all-RNA oligonucleotide to a DNA oligonucleotide with five ribonucleotides at its 3′ terminus (i.e., nearest the amino acid) ameliorated the issue for both Cys and Asn, suggesting that the amino acid-proximal ribonucleotides of the 5′ adapter interfere with ligation, and by extension, sequencing of Cys and Asn bridged tRNAs (**Fig. S2C,D**).

### Discrimination of charged and uncharged biological tRNAs by nanopore sequencing

Modified basecalling of nanopore sequencing signals is achieved by training neural networks (*39*); in the case of direct RNA sequencing, detecting the presence of a modification via comparison to an unmodified control is often a more trivial computational challenge than precise characterization of a modification’s identity (*40–43*). To extend this concept to aminoacylated tRNA sequencing, we sought to classify charged and uncharged tRNA sequencing reads, with the goal of developing a one-pot approach for nanopore tRNA sequencing to capture charged and uncharged molecules within the same sequencing library. We prepared separate “ground truth” libraries containing either (***i***) chemically ligated, charged tRNA, or (***ii***) deacylated, enzymatically ligated tRNA from budding yeast (**Fig. 2A**). tRNAs were ligated to 3′ adapters containing unique sequences for charged and uncharged molecules, and a common 5′ adapter incorporating the design changes (DNA/RNA hybrid) from in **Fig. S2B**, and sequenced these libraries on PromethION *RNA* flow cells.

**Figure 2.**
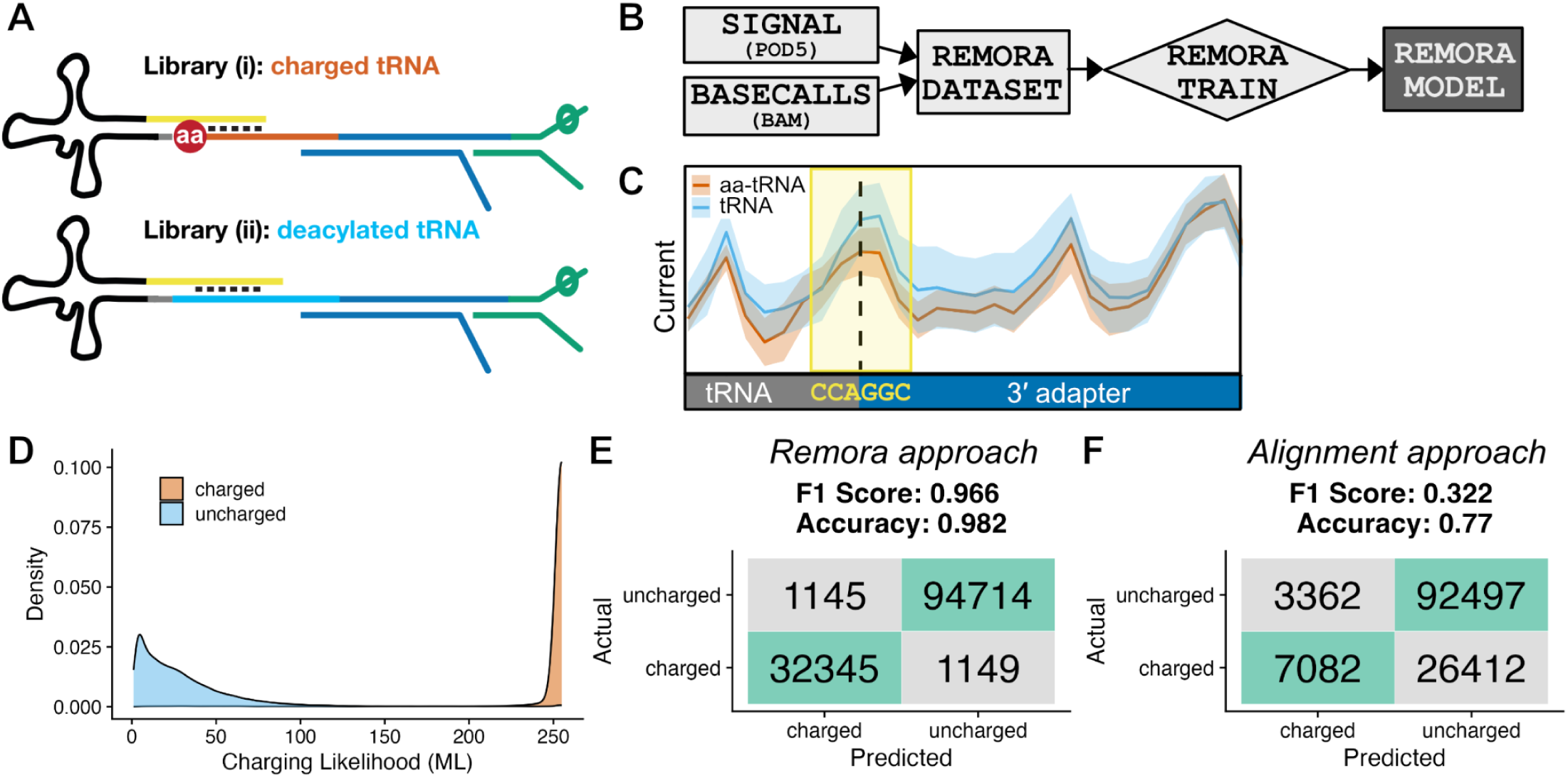
Model training and classification of aminoacylated tRNA reads from nanopore signal. (**A**) Schematic illustrating ground truth libraries for model training. Library *(i)* was prepared by only chemical ligation to biological aminoacylated tRNAs. Library *(ii) was* prepared by only enzymatic ligation to deacylated biological tRNAs. (**B**) Strategy for dataset preparation and neural network training of a Remora model to identify charged and uncharged tRNA reads from nanopore current signals. (**C**) Schematic depicting reference-anchored signal from libraries *(i) and (ii)*, with the 6-nt training window enclosed within the yellow box, and the position of the amino acid via dashed line. Mean current signals in picoamps for charged and uncharged tRNA reads are indicated by solid lines within each colored trace, with corresponding shading spanning the standard deviation. (**D**) Density plot showing the distribution of aminoacylation prediction scores (modification likelihood or “ML” score, 0-255 scale) assigned by Remora model in validation libraries (95,859 deacylated tRNA reads, 33,494 aminoacylated tRNA reads). (**E**) Confusion matrix and key metrics illustrating the model’s performance at classifying a second set of replicates prepared as in (**A**), using a cutoff of ML > 199 for identification of aminoacylated reads from nanopore signal. (**F**) Confusion matrix when a sequence alignment approach was used for charged tRNA classification on the same validation dataset.

Informed by our synthetic tRNA sequencing results (**Fig. 1F**), we trained a convolutional neural network on this dataset using the Remora software from ONT (**Fig. 2B**), which predicts the presence of modified nucleotides using ionic current signals generated during nanopore sequencing. For training, we defined a six-nucleotide signal window spanning the invariant CCA at the 3′ terminus of mature tRNA (“CCA-G-GC”, where “G” represents the first base of adapter most affected by the amino acid, and “GC” represent the next two bases of the adapter; depicted in **Fig. 2C**), as the sequence in this region would be shared across all tRNA isodecoders. We generated a Remora dataset from 80% of these reads, with labels defined by the library of origin (***i*** and ***ii***, above), and then trained and evaluated a model on the reserved 20% test set. During signal-anchored inference, Remora models output predictions for “modified” positions in the ML (“modification likelihood”) tag of a BAM file; these values range from 0-255, where lower values are more likely to be canonical nucleotides. Inspection of these values in our libraries revealed a bimodal distribution, with 96.6% of reads from the charged library (***i***) bearing ML scores ≥200, compared to 1.2% of in the deacylated sample (***ii***) (**Fig. 2D**).

We generated a second biological replicate of the libraries in **Fig. 2A** for model validation, performing reference-anchored inference using the trained model above and using a ML cutoff of >199 to classify aminoacylated tRNAs. **Fig. 2E** illustrates the Remora model’s performance on this new dataset, with an F1 score of 0.966. This signal-based approach substantially outperformed an alignment-based one using unique 3′ adapter sequences to discriminate charged and uncharged tRNAs, which yielded an F1 score of 0.322 with low sensitivity for identifying charged tRNA reads (**Fig. 2F**), and which we attribute to increased base-calling error caused by the embedded amino acid. Based on its clear improvement over the alignment-based approach, we focused on signal-based classification of tRNA aminoacylation state.

### Measurement of tRNA aminoacylation with aa-tRNA-seq

To determine whether this approach could report on tRNA charging levels, we generated replicate aa-tRNA-seq libraries from wild-type budding yeast under conditions identical to **Fig. 1B** (wild-type yeast in rich media), enabling an orthogonal comparison of tRNA aminoacylation measured by acidic northern (**Fig. 1B**) and model-based classification of nanopore sequencing reads. Across the 16 tRNA isodecoders quantifiable by densitometry (**Fig. S1D**), we observed a Pearson’s correlation of 0.68; however, estimates of isodecoder charging by sequencing were uniformly lower than charging levels measured by northern (**Fig. 3A**). While in principle this difference might be explained by differences in the rates of aberrant read termination between charged and uncharged molecules, the magnitude of this result was inconsistent with our observations in synthetic tRNA sequencing data after optimization of the 5′ adapter sequence (**Fig. S2**). We therefore re-analyzed the initial dataset used for Remora model training, comparing the translocation time for tRNA reads in the first library prepared via chemical ligation to aminoacylated tRNA to the second library where tRNAs were deacylated and enzymatically ligated. This analysis revealed that the total translocation time for aminoacylated tRNA—across all isodecoders—is on average ∼1.2-fold longer than their uncharged counterparts (**Fig. S3**), which is likely the primary driver of the underestimation of charging we observe in aa-tRNA-seq libraries (**Fig. 3A**): in a complex mixture of biological tRNAs, uncharged tRNAs are sequenced faster than charged tRNAs, leading to a sampling bias toward uncharged tRNAs.

**Figure 3.**
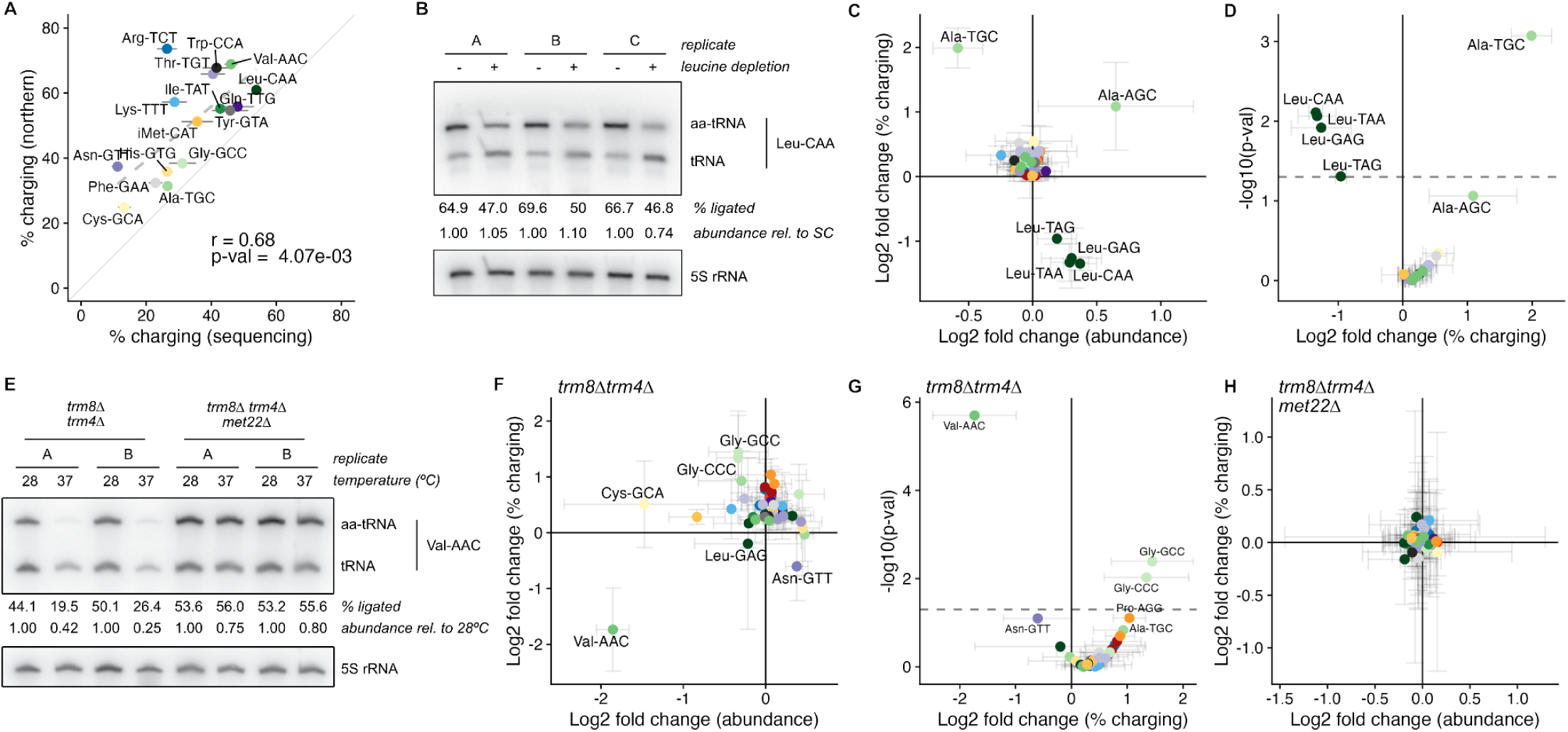
Sequencing and analysis of budding yeast tRNAs via chemical-charging northern and aa-tRNA-seq. (**A**) Correlation of tRNA aminoacylation (% charging) in budding yeast, as measured by acid northern in **Fig. 1B** (y-axis), and aa-tRNA-seq (mean and standard deviation of 3 replicates; dashed gray Pearson correlation with r and p-value; solid gray *y=x* line). (**B**) Chemical-charging northern analysis of Leu-CAA aminoacylation in a budding yeast leucine auxotroph upon 15 minutes of leucine depletion, as measured by the percent of tRNA chemically ligated in each lane (upper band), with the percent-ligated tRNA per sample quantified under each lane. Signals were normalized to 5S rRNA probe (lower inset); relative abundances represent within-replicate normalized levels of total tRNA (with the sample grown in complete media normalized to 1.0 and compared to the abundance for each leucine-starved sample). (**C**) Log_2_ fold change in tRNA abundance and tRNA charging percent charged reads for all isodecoders after 15 minutes growth in synthetic complete (SC) or leucine dropout media. Points represent the mean of the same 3 biological replicates from panel (**B**), with error bars spanning the standard deviation. (**D**) Volcano plot of the mean fold change in aminoacylation for the same 3 replicates in panels (**C**) and (**D**), with Z-test p-values on the y-axis and dashed line indicating the *α* threshold. (**E**) Chemical-charging northern analysis of chemically ligated tRNA from two biological replicates of the RTD-sensitive budding yeast strain *trm8Δ trm4Δ* and the RTD-resistant strain *trm8Δ trm4Δ met22Δ* grown at permissive (28°C) and nonpermissive (37°C) temperatures for three hours. The percent of chemically ligated Val-AAC tRNA (upper band) is indicated below each lane, with a 5S rRNA probe as a loading control in the lower inset. The relative abundances represent within-replicate normalized levels of total tRNA (with the samples grown at 28°C normalized to 1.0 and compared to the abundance for a matched sample shifted to 37°C). (**F**) Log_2_ fold-change in tRNA abundance and tRNA charging percent charged reads across 3 biological replicates upon shift to nonpermissive temperature in *trm8Δ trm4Δ* cells. (**G**) Volcano plot of the mean fold change in aminoacylation from panel (**F**), with Z-test p-values on the Y axis and dashed line indicating the *α* threshold. (**H**) Log_2_ fold-change in tRNA abundance and tRNA charging percent charged reads across 3 biological replicates upon shift to nonpermissive temperature in *trm8Δ trm4Δ met22Δ* cells.

### aa-tRNA-seq detects changes in tRNA aminoacylation during nutrient stress

We next sought to assess whether aa-tRNA-seq could detect dynamic changes in tRNA charging and focused on nutrient limitation, which causes rapid changes to the aminoacyl-tRNA pool. We subjected a *S. cerevisiae* leucine auxotroph to leucine starvation for 15 minutes, and found that, as previously reported (*11*), this caused rapid depletion of aminoacylated leucyl-tRNA isodecoders (**Fig. 3B-D**). Using a chemical-charging northern (**Fig. 3B**), we observed a mean 28.4% decrease in the abundance of aminoacylated leucyl-tRNA. We performed aa-tRNA-seq on the same material, and examined global changes in tRNA abundance and aminoacylation upon leucine starvation. This analysis revealed a significant decrease in the charging of all four leucyl tRNA isodecoders present in budding yeast, while most other tRNAs remained unaffected. Surprisingly, we also found a concomitant increase in the charging of Ala-tRNA isodecoders (3.97-fold for Ala-TGC, p < 0.001; 2.12-fold for Ala-AGC, p = 0.08; BH-adjusted Z test; **Fig. 3C,D**), an effect that was not detected in earlier microarray-based experiments examining tRNA charging in response to leucine starvation (*11*).

### aa-tRNA-seq detects interdependence between tRNA modifications and aminoacylation during rapid tRNA decay

Hypomodified tRNAs are susceptible to “rapid tRNA decay” (RTD) in budding yeast, fission yeast, and bacteria (*44–52*). For example, Val-AAC tRNA lacking 5-methyl cytidine and 7-methyl guanine in *trm4Δ trm8Δ* budding yeast is rapidly destabilized and degraded at high temperature by 5′-3′ exonucleolytic decay (*53*). We cultured a *trm4Δ trm8Δ* strain and a control bearing an additional disruption of *MET22* at the permissive temperature of 28°C followed by a shift to the non-permissive temperature of 37°C for three hours. Deletion of *MET22* causes accumulation of pAp (adenosine 3′,5′ bisphosphate, a competitive inhibitor of 5′-3′ exonucleases (*54*)), suppressing RTD (*53*). We isolated small RNA from each strain and temperature in biological triplicate, and prepared this material for analysis by chemical-charging northern and aa-tRNA-seq. Both approaches confirmed defects in both stability and aminoacylation for Val-AAC (*53*). By chemical-charging northern, aminoacylation levels for Val-AAC drop approximately 2-fold upon shift to the nonpermissive temperature in *trm8Δ trm4Δ* cells, and this effect was suppressed by *met22Δ* (**Fig. 3E**). This effect is readily detected by aa-tRNA-seq, where Val-AAC undergoes significant changes in both tRNA abundance and aminoacylation upon temperature shift (**Fig. 3F**). We also observed statistically significant increases in aminoacylation (2.54-fold for Gly-GCC, p-value = 0.009; 2.73-fold for Gly-CCC, p-value = 0.004, BH-adjusted Z test) for two glycyl-tRNA isodecoders upon a shift to 37°C (**Fig. 3F,G**), which were eliminated by *met22Δ* (**Fig. 3H**).

### A machine learning approach to discriminate embedded amino acids in aa-tRNA-seq

Because the identity of the amino acid on an aminoacylated tRNA is invisible to current tRNA sequencing approaches, a major advance for the field would be the detection of misaminoacylation events directly from high throughput tRNA sequencing libraries. Towards this goal, we performed detailed analysis of signals produced during nanopore sequencing for aminoacyl-tRNA charged with each of the 20 proteinogenic amino acids using the Flexizyme system. While nanopore basecallers and other machine learning approaches for detection of modified bases are typically trained on ionic current signatures produced at specific residues (*43*, *55*, *56*), the raw signal generated during nanopore sequencing is a composite of changes in current and translocation speed (measured in “dwell time”, the time between inferred translocation states). Because they are sequence-identical (with the exception of Cys- and Asn-tRNA libraries, which were prepared using the DNA/RNA 5′ adapter sequence tested in **Fig. S2** and used in all biological sequencing experiments), our synthetic aa-tRNA-seq libraries enable the isolation of amino acid specific signals by comparing each of the 20 aa-tRNAs to an uncharged control.

Aminoacylated tRNAs yield specific distortions in nanopore signals, generating unique signatures that vary between amino acids. **Fig. 4A** displays the mean dwell time in milliseconds for each of our 20 synthetic aa-tRNA libraries and an uncharged control tRNA. We observed large increases in dwell time for charged tRNAs at a position located 9 nucleotides downstream from the amino acid (position 86), with dwell times exceeding 1 second for 9 of 20 amino acids. Because nanopore direct RNA sequencing proceeds in a 3′ to 5′ direction, we speculate that this signal represents specific interactions between amino acid and the motor protein as the 3′ adapter is transiting through the nanopore reader head. **Fig. 4B** shows the relative change in normalized current for each of the aminoacylated tRNA reads over this same window, compared to the uncharged tRNA control. Charged tRNA libraries display lower mean current values than the non-aminoacylated substrate across most of this region, suggesting that amino acids occlude ionic flow as they transit through the helicase/pore assembly. Consistent with this explanation, we observed the largest reductions in mean current at the precise site of aminoacylation for the most bulky amino acids, indicating that the largest distortions in ionic current distortions occur within the narrowest aperture of the nanopore itself.

**Figure 4.**
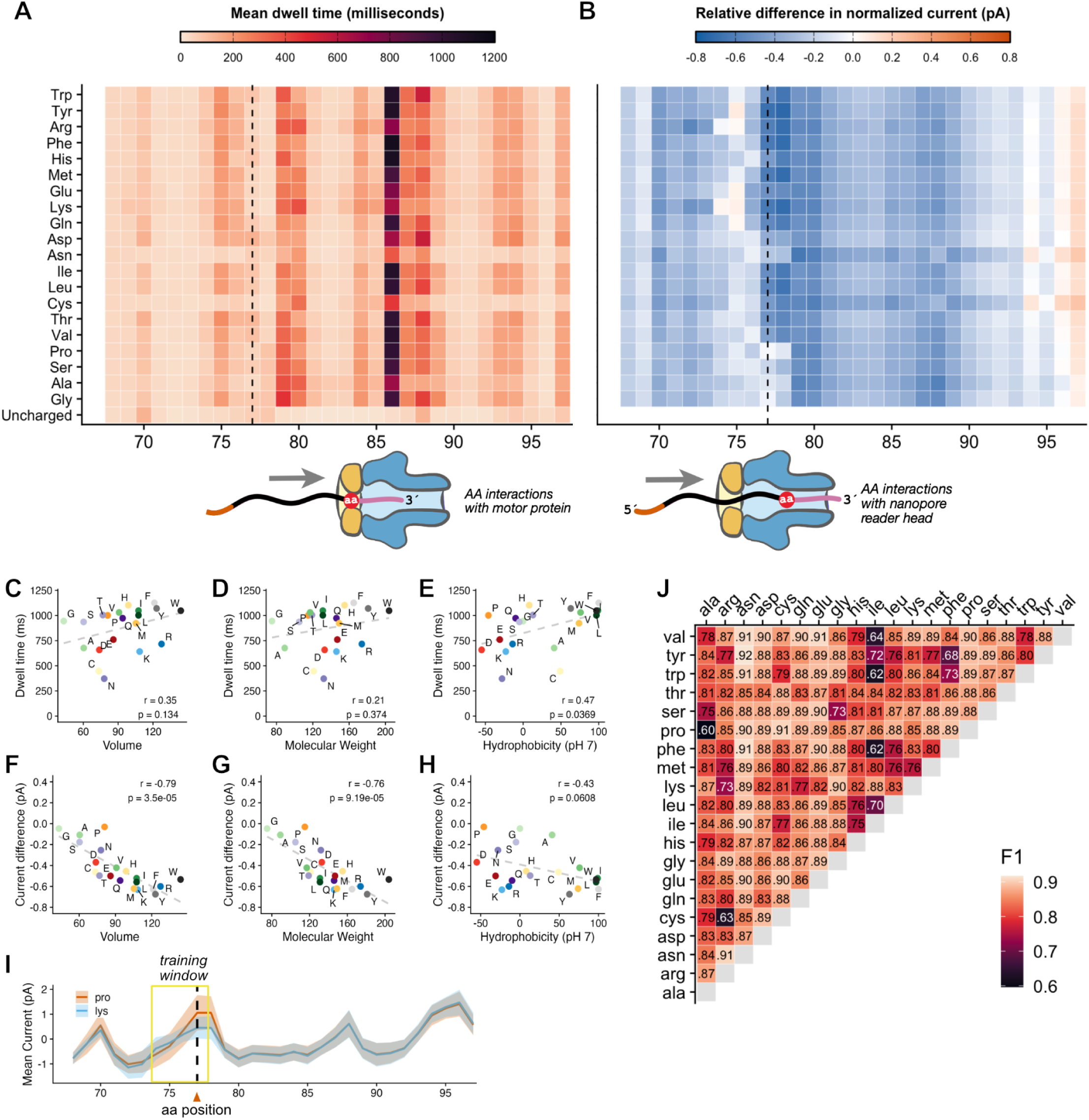
Signal analysis and classification of amino acid identity using nanopore sequencing. (**A**) Mean dwell time for a synthetic tRNA aminoacylated with 20 naturally-occurring amino acids as well as an uncharged control that has been enzymatically ligated to the 3′ adapter prior to nanopore direct RNA sequencing. Sequences are ordered top to bottom on the y-axis by amino acid side chain molecular weight. The plotted region includes the 3′ terminus of the tRNA (6 nt, positions 68-73), the CCA tail (positions 74-76), the aminoacylated position (dashed line, position 77) and the entirety of the 3′ adapter sequence. The schematic underneath the heatmap depicts the direction of translocation and approximate location of the amino acid within the helicase motor protein (in orange) when the 3′ adapter sequence is centered in the nanopore (blue) and the largest increases in dwell time are observed. (**B**) Relative differences in normalized mean current between for each synthetic aminoacylated tRNA compared to the uncharged control, over the same region as in panel (**A**). The lower schematic illustrates the location of the amino acid within the nanopore reader head when the largest changes in current are observed. (**C**) Scatter plot with linear regression line showing the relationship between mean dwell time at position 86 and amino acid volume. Points represent individual amino acids, with side-chain volume (in cubic angstroms) on the x-axis and the relative difference in mean current on the y-axis. A dashed line indicates the linear regression fit, with the Pearson’s correlation coefficient (r) and p-value displayed. (**D**) Scatter plot showing the relationship between mean dwell time at position 86 and amino acid molecular weight (grams / mole). (E) Scatter plot showing the relationship between mean dwell time at position 86 and the hydrophobicity index (*80*) for each amino acid at pH 7, with higher values being more hydrophobic. (**F**) Scatter plot showing the relationship between mean current differences at the aminoacylated position (position 77, dashed line in **C**) and amino acid volume (cubic angstroms). (**G**) Scatter plot showing the relationship between mean current differences at the aminoacylated position (position 77, dashed line in **C**) and amino acid molecular weight (grams / mole). (**H**) Scatter plot showing the relationship between mean current differences at the aminoacylated position (position 77, dashed line in **C**) and the hydrophobicity index for each amino acid. (**I**) Schematic depicting pairwise Remora model training strategy for differentiation of reads containing individual amino acids, using distribution of current signals for synthetic tRNA charged with proline vs lysine as an illustrative example. The mean current in picoamps for the lysyl-tRNA substrate is represented by the central blue line, the shaded blue area representing the standard deviation around the mean, and tRNA-Pro data plotted analogously in red. A yellow box outlines the window of signal (4 nt, with one representing a placeholder for the amino acid) on which each pairwise model was trained, with the location of the chemically ligated amino acid indicated by the dashed line. (**J**) F1 scores for all pairwise Remora models trained on the 20 synthetic tRNA substrates using the strategy illustrated in **(I)**.

We examined the correlation between the largest current and dwell time effects and various amino acid properties. While the putative helicase interactions at position 86 are poorly correlated with amino acid volume and molecular weight (**Fig. 4C,D**), we identified strong correlations between dwell time and hydrophobicity (**FIg. 4E**), as well as correlations between amino acid mass and volume, and the observed current differences at position 77 (the position of the amino acid; **Fig. 4F,G**). Together, these observations likely reflect straightforward physical occlusion of ion flow within the nanopore reader head and more complex interactions between the tRNA-embedded amino acid and the motor protein.

Closer examination of our data revealed substantial variation in the magnitude of current (**Fig. S4**) and dwell time (**Fig. S5**) signals produced by different synthetic aminoacyl-tRNAs. We trained 380 pairwise models to discriminate one AA from another (**Fig. 4I)**, and evaluated their performance on our synthetic data (**Fig. 4J**). The performance of these models was generally high with a median F1 score of 0.86, indicating that differences in nanopore current between several pairs of amino acids represent a strong signal for classification. Notable underperforming outliers included isoleucyl-tRNAs, which are poorly distinguished from other hydrophobic side chains, and other specific comparisons (e.g., Phe-Tyr, Pro-Ala, Cys-Arg). We are currently exploring the improvement of these models by explicit inclusion of dwell time information during model training, and plan to apply these models to understand specific cases of potential tRNA misacylation (**Fig. 3D,G**).

## DISCUSSION

We developed a novel chemical ligation approach to selectively capture aminoacylated tRNAs, and used it to study tRNA aminoacylation in a variety of contexts. First, we leveraged chemical ligation to simplify the analysis of aminoacylated tRNAs via northern blotting (**Fig. 1C**). This approach offers key advantages, including simplified electrophoretic separation of chemically ligated species, and is therefore a compelling alternative to more laborious techniques. However, it is important to note that under the existing reaction conditions, there is some variability in ligation efficiency among biological tRNA isoacceptors and only some were ligated quantitatively (**Fig. S1C,D**), limiting the performance of this method compared to a traditional acidic northern blot. It is also noteworthy that chemical ligation is likely useful beyond the capture of aminoacylated tRNA: a variety of nucleophiles can react with phosphorimidazole-activated oligonucleotides, including RNA terminal hydroxyl groups at elevated pH (*57*). Juxtaposition of the phosphorimidazolide near the ɑ-amino group of the aa-tRNA was facilitated by splinted base-pairing to the 3′-CCA overhang (**Fig. 1A**), but one could imagine other sequence-specific targeting scenarios, including the 3′-ends of mRNA poly-A tails.

Next, we adapted the chemical ligation to enable direct nanopore sequencing of aminoacylated tRNA (“aa-tRNA-seq”). We first trained a Remora classification model to distinguish charged and uncharged tRNAs from nanopore sequencing data, and showed that this signal-based classification approach outperformed an alignment-based one (**Fig. 2**). Our application of aa-tRNA-seq confirmed known effects of hypomodification and nutrient deprivation on tRNA stability and aminoacylation (**Fig. 3**). Our analysis of RTD mutants demonstrates that aa-tRNA-seq can identify interactions between tRNA hypomodification and aminoacylation, and revealed the effects of *TRM8* and *TRM4* disruption on tRNA turnover to be uniquely specific to Val-AAC (**Fig. 3F**). We also found significantly increased aminoacylation for two isoacceptor families in cases of hypomodification (Gly-GCC and Gly-CCC, **Fig. 3G**) and nutrient deprivation (Ala-TGC, **Fig. 3D**). These may simply be increases in cognate acylation of these isodecoders that were not previously observed. Alternatively, they may represent new cases wherein a lack of modification or charging enables misaminoacylation of Gly and Ala isodecoders. Methionine misaminoacylation is a common response to stress (*58*) wherein the methionyl tRNA synthetase non-specifically charges several tRNA isodecoders with methionine. However, the pattern of misaminoacylation we observe is restricted to a few tRNAs, suggesting an alternative mechanism.

Like other tRNA sequencing methods, aa-tRNA-seq has tradeoffs and technical biases. First, we identified intrinsic differences in the total translocation time between charged and uncharged tRNA during nanopore sequencing (**Fig. S3**), which is a barrier to absolute quantitation of tRNA charging. Second, under these reaction conditions not all biological tRNA isoacceptors are quantitatively ligated (**Fig. S1C**), reflecting an additional source of bias that may be further optimized. This issue of ligation efficiency also reflects a broader challenge: enzymatic adapter ligation bias has complicated other tRNA sequencing methods, often making comparisons of tRNA abundances within the same sample unreliable (*59*, *60*). We note that while methods relying on periodate oxidation to distinguish charged from uncharged tRNAs have proved valuable to the field, they also face technical challenges, including tRNA damage during periodate treatment (*21*), variability in the sensitivity of tRNA 3′ termini to oxidation, and inconsistency in the number of nucleotides removed during treatment (*61*). Additionally, as all published periodate-based methods require cDNA generation, they are subject to biases from tRNA modifications that inhibit reverse transcriptase processivity. Alternatively, if these modifications are instead removed, such methods can introduce additional RNA damage from harsh buffer conditions during enzymatic pretreatment (*62–64*).

Chemically ligated aa-tRNAs generate discrete signals during nanopore sequencing due to interactions between the embedded amino acid and the motor protein and nanopore architecture (**Fig. 4**). This unique feature enabled accurate identification of *bona fide* aa-tRNAs while minimizing false positives: we classified 3.6% of reads from the "charged-only" validation library as uncharged. However, some of these may be true non-acylated tRNA that were enzymatically ligated to hydrolyzed phosphorimidazole-adapter, which has a 5′-phosphate terminus competent for T4 RNL2 ligation. While our current approach demonstrates that different amino acids produce distinct signal properties that enable their discrimination in pairwise comparisons (**Fig. 4J**), further advances in our machine learning approach may enable the direct, *de novo* identification of amino acids in biological samples. Such improvements would enable the reanalysis of existing aa-tRNA-seq data, adding new power to this approach and unlocking new questions about tRNA charging and misaminoacylation. We also anticipate that incorporation of translocation time information into models for classifying aminoacylated tRNAs will generate additional improvements to aa-tRNA-seq. Dwell time provided useful information in the detection of RNA modifications using the previous direct RNA (RNA002) ONT chemistry, including pseudouridine (*65*), 2′-O-methyl (*66*), and 2′-phosphate (*31*) modifications, but existing approaches for training models on nanopore signal do not leverage this information directly, due to the fact that Remora and other software re-anchor ionic current information onto an aligned sequence. While dwell time information is retained in this process, it is transformed and thereby de-emphasized in model training. Notably, the increased dwell times observed for aminoacylated tRNAs—likely due to unique helicase interactions—are a robust signal in our data. However, these effects are less strongly correlated with amino acid properties than the current differences at the aminoacylated position (**Fig. 4C-H**), where we have focused our model training.

The translocation rate for aa-tRNAs also impacts the pore blocking effects we described for Cys- and Asn-tRNA (**Fig. S2**), as read ejection (“unblocking”) is initiated during nanopore sequencing when a constant signal (indicating a stalled molecule) exceeds a set time threshold. While we successfully resolved pore blocking issues via optimization of the 5′ adapter sequence, we do not fully understand why chemical ligation of these substrates produced these artifacts, nor why they were resolved by the substitution of deoxyribonucleotides in our 5′ RNA/DNA splint adapter. While Asn-tRNA yielded significant pore blocking, Gln-tRNA did not (**Fig. S2A**), suggesting an issue beyond simply the presence of an amide side chain. We do not fully understand this difference, but note that the Asn side is uniquely capable of cyclization rearrangements during intein catalysis, which may contribute to its unique pore blocking phenotype (*67*).

Transfer RNA modifications are installed in evolutionarily conserved but incompletely understood circuits (*68*, *69*). While multiple links between tRNA modifications and aminoacylation have been reported (*4*, *5*), these examples remain sparse and poorly characterized, due in part to the lack of incisive and accessible tools to study these relationships. Comprehensive identification and characterization of circuits linking modification and aminoacylation along with tRNA abundance will require further optimization not only of the approaches outlined in this manuscript, but also the ongoing work of mapping the 67+ unique RNA modifications present in the tRNA epitranscriptome (*70*), which will likely require a combination of nanopore sequencing, mass spectrometry (*71*, *72*), and other technologies, with the aim of characterizing both the complete collection of tRNA modifications (using lower throughput approaches) and understanding the signals they produce both singly and in combination during high throughput, direct RNA sequencing experiments (*73*). Moreover, increased accuracy of aa-tRNA identification may further enhance our understanding of how tRNA modifications act as determinants or anti-determinants of synthetase recognition and specificity. Moreover, direct identification of amino acids and their corresponding tRNA sequences could advance studies on aminoacyl-tRNA synthetase evolution and engineering, bypassing the indirect readouts and negative selections commonly employed by current synthetase engineering approaches (*74*, *75*).

Transfer RNAs are long-lived and undergo repeated cycles of charging and deacylation throughout their lifetime. While tRNA biogenesis is recognized as an intricately coordinated process—spanning transcription, processing, modification, and aminoacylation—less is known about the molecular transformations that tRNAs undergo throughout their functional lifetime, or the specific events that trigger their turnover. Because tRNAs undergo continuous transformation and are highly responsive to cellular conditions, we anticipate that application of this method will yield novel insights into how cells adapt to stress and disease.

## MATERIALS AND METHODS

### Preparation and chemical ligation of synthetic aminoacylated tRNAs

#### Synthesis

Oligonucleotides listed in **Table 2** were either purchased from IDT or synthesized on the K&A H-6 RNA/DNA synthesizer. Phosphoramidite coupling times and the remaining synthesis method parameters were as instructed by the manufacturer (ChemGenes and Glen Research). After solid-state synthesis, oligonucleotides were cleaved and the nucleobases deprotected as recommended by ChemGenes and Glen Research. The cleaved and deprotected solutions were evaporated using a speed-vac for 2 hours followed by overnight lyophilization. The dry material was dissolved in 100 µL DMSO to which 125 µL of TEA. 3HF was added followed by incubation at 65°C for 2.5 hours. The fully deprotected oligonucleotides were precipitated with 0.1 volumes of 5 M ammonium acetate and 5 volumes of cold isopropanol. The precipitated material was dissolved in 5 mM EDTA in 99 % v/v formamide and purified by denaturing PAGE. The desired gel bands were visualized by UV shadowing, cut out, crushed, and soaked in 2 mM EDTA, 5 mM sodium acetate on a rotator overnight. The rotated solutions were filtered through a 5 µm syringe filter after which the filtered solutions were concentrated using Amicon MWCO filters. The concentrated solutions were finally precipitated using 0.1 volumes of 3 M sodium acetate and 5 volumes of ethanol, washed twice with 80 % v/v ethanol, and air dried.

The 3,5-dinitrobenzyl esters of amino acids (DBE-aas) were synthesized as described in ref. 34 with the following modifications:

1. Boc protecting groups were removed by dissolving the dry crude DBE-Boc-aa material in 2 mL neat TFA and incubating it at room temperature for 10 minutes. The TFA was removed under a stream of nitrogen and the deprotected product was washed twice with diethyl ether. The diethyl ether was removed under vacuum and the final DBE-aa product was dissolved in DMSO and used in the aminoacylation assays.
2. Boc-Ser, Boc-Thr, and Boc-Tyr (ChemImpex) were purchased with O-*tert*-butyl protection on the side chain. The deprotection was performed in 90:10 TFA:triethylsilane for 2 hours.
3. Boc-Met (ChemImpex) was used without additional side chain protection, but during the TFA deprotection two side products were observed: oxidation to produce a DBE-Met dimer and *tert*-butylation of the sulfur. This necessitated reversed-phase purification. RediSep Gold® C18 Reversed Phase Column was used with the 5-90 % gradient of solvent B (solvent A = 2 mM TEAB pH 8; solvent B = acetonitrile).
4. Boc-Gln and Boc-Asn were purchased with Xan protection on the side chain amide (ChemImpex). DBE-Asn required reversed-phase purification. RediSep Gold® C18 Reversed Phase Column was used with the 5-90 % gradient of solvent B (solvent A = 2 mM TEAB pH 8; solvent B = acetonitrile). DBE-Asn additionally requires immediate use after purification due to the presumed rapid intramolecular attack of the side chain amide onto the activated ester.
5. Boc-Cys was purchased with Trt protection on the side chain thiol (ChemImpex). The deprotection was performed in 90:10 TFA:triethylsilane for 2 hours.

#### Activation

The 5′-phosphorimidazolide adapter was generated by incubating a solution containing 200 µM of the 5′-phosphorylated adapter, 100 mM imidazole pH 7, and 100 mM EDC.HCl for 2 hours at room temperature. The activated adapter was then precipitated by adding 0.1 volumes of saturated sodium perchlorate in acetone and 3 volumes of cold acetone. The pellet was washed twice with a 1:1 v/v solution of acetone:diethyl ether followed by drying under vacuum. The activated adapter was dissolved in 1 mM imidazole pH 8 and stored at −80°C until use. The same stock of the activated adapter was used throughout the experiment, but care was taken to thaw the stock immediately prior to the experiment, to store it on ice while using it, and to return it to −80°C as quickly as possible.

#### Aminoacylation

The aminoacylation reactions containing 100 mM HEPES pH 8, 10 mM MgCl_2_, 40 µM synthetic tRNA, 36.7 µM dFx Flexizyme, 5 mM DBE-aa (20 % v/v DMSO), were incubated on ice for 16 hours.

#### Chemical ligation

The ligation reactions were set up by diluting the aminoacylation reactions 8-fold so that the final solution contained 5 µM of the aminoacylated tRNA, 50 µM 5′-phosphorimidazolide adapter, 50 µM splint (also the 5′-adapter below), 5 mM EDTA, and 37.5 mM HEPES pH 8. The reactions were allowed to proceed for 24 hours on ice, before being diluted with an equal volume of a solution of 5 mM EDTA in 99 % v/v formamide and purified by 16 % denaturing PAGE. The ligated products were cut out from the gel, crushed, and soaked in a solution of 2 mM EDTA, 5 mM sodium acetate, acidified to pH 5 on a rotator for 3 hours at 4 °C. The extracted aa-bridged tRNA products were then filtered using 0.22 µm spin filters, concentrated using Amicon 10k MWCO filters, and desalted using the Oligo Clean and Concentrator kit (Zymo Research).

#### Cys-bridged tRNA alkylation

After chemical ligation and gel purification, the Cys-bridged tRNA was reduced with DTT for 1 hour at room temperature. The reaction contained 1.2 µM Cys-bridged tRNA, 50 mM HEPES pH 8, and 10 mM DTT. After the 1 hour incubation, the reduction reaction was diluted 1.33-fold so that the final alkylation solution contained 0.9 µM Cys-bridged tRNA, 37.5 mM HEPES pH 8, 7.5 mM DTT, and 50 mM chloroacetamide. The alkylation reaction was allowed to proceed for 30 mins in the dark, after which it was cleaned up using the RNAClean XP beads (Beckman Coulter) according to the manufacturer protocol with the following change: immediately after the addition of the bead suspension to the ligation reaction isopropanol equal to volume of the reaction+beads was added.

### Nanopore library preparation and sequencing of synthetic aminoacylated tRNAs

The chemically ligated tRNA products from above were enzymatically ligated to the 5′-adapter/splint for 30 minutes at room temperature. The ligation reactions contained 16 pmol of the chemically ligated tRNA, 80 pmol of the 5′-adapter, 1x NEB T4 RNA Ligase 2 buffer supplemented with 5% PEG 8000, 2 mM ATP, 6.25 mM DTT, 6.25 mM MgCl2, and 0.5 units/μL T4 RNA ligase 2 (10,000 units/mL). The ligated material was purified using the RNAClean XP beads (Beckman Coulter) as above. This material was then prepared for nanopore direct RNA sequencing via RTA ligation, which was performed using tRNA purification specific magnetic beads (BioDynami Cat.# 40054S). The remaining library prep and nanopore sequencing was performed as described below on P2solo sequencing instruments, using MinKNOW version 23.11.7.

### Synthetic tRNA mini-substrate experiments

The acceptor stem mimic oligonucleotide (**Table 2**) was aminoacylated with all 20 amino acids as described above. At the end of the 16 hour incubation:

1. 1 µL aliquots were diluted in 9 µL of acidic quenching buffer (10 mM EDTA pH 8.0, 1x bromophenol blue, 100 mM sodium acetate pH 5.0, 150 mM HCl, 75 % v/v formamide) and analyzed by 20 % acidic denaturing PAGE (acidic gels contained 100 mM sodium acetate pH 5.0 instead of the usual 1x Tris-Borate-EDTA). The acidic gels were run in 100 mM sodium acetate pH 5 at 25 W for 3 hours at 4 °C. Aminoacylation percentage was obtained by quantifying the per-lane normalized band intensity in the ImageQuant TL software. The amino acids that displayed sufficient aminoacylated versus nonaminoacylated gel band resolution were subjected to the next step.
2. The remaining aminoacylation reaction was immediately diluted 10-fold in the chemical ligation buffer in three separate replicates. The ligation reaction contained 1 µM of the aminoacylated RNA, 4 µM of the 5′-phosphorimidazolide activated hairpin adapter (**Table 2**), 10 µM of the acceptor stem mimic complement (**Table 2**), 200 mM HEPES pH 6.5, 5 mM MgCl_2_, and 100 mM of 1-(2-Hydroxyethyl)imidazole pH 6.5. After 90 mins at room temperature, 1 µL aliquots were diluted in 9 µL of acidic quenching buffer, and analyzed by standard 20 % denaturing urea-PAGE. The efficiency of the ligation reaction was obtained by quantifying the per-lane normalized band intensity in the ImageQuant TL software. The normalized ligation efficiency was obtained by dividing the fraction ligated by the fraction aminoacylated and multiplying by 100 %.

### Yeast strains and growth conditions

Yeast strains used in this study are listed in **Table 1**. For chemical ligation validation by acid northern (**Fig. 1B**), a single colony of S288C was inoculated into YEP glucose (yeast extract, peptone, 2% glucose) and incubated at 30°C overnight with rotation before dilution to an OD_660_ of 0.2 in YEPD media. The culture was grown to log phase shaking at 30°C before a pellet was collected, flash frozen in liquid nitrogen, and stored at −80°C.

**Table 1.**
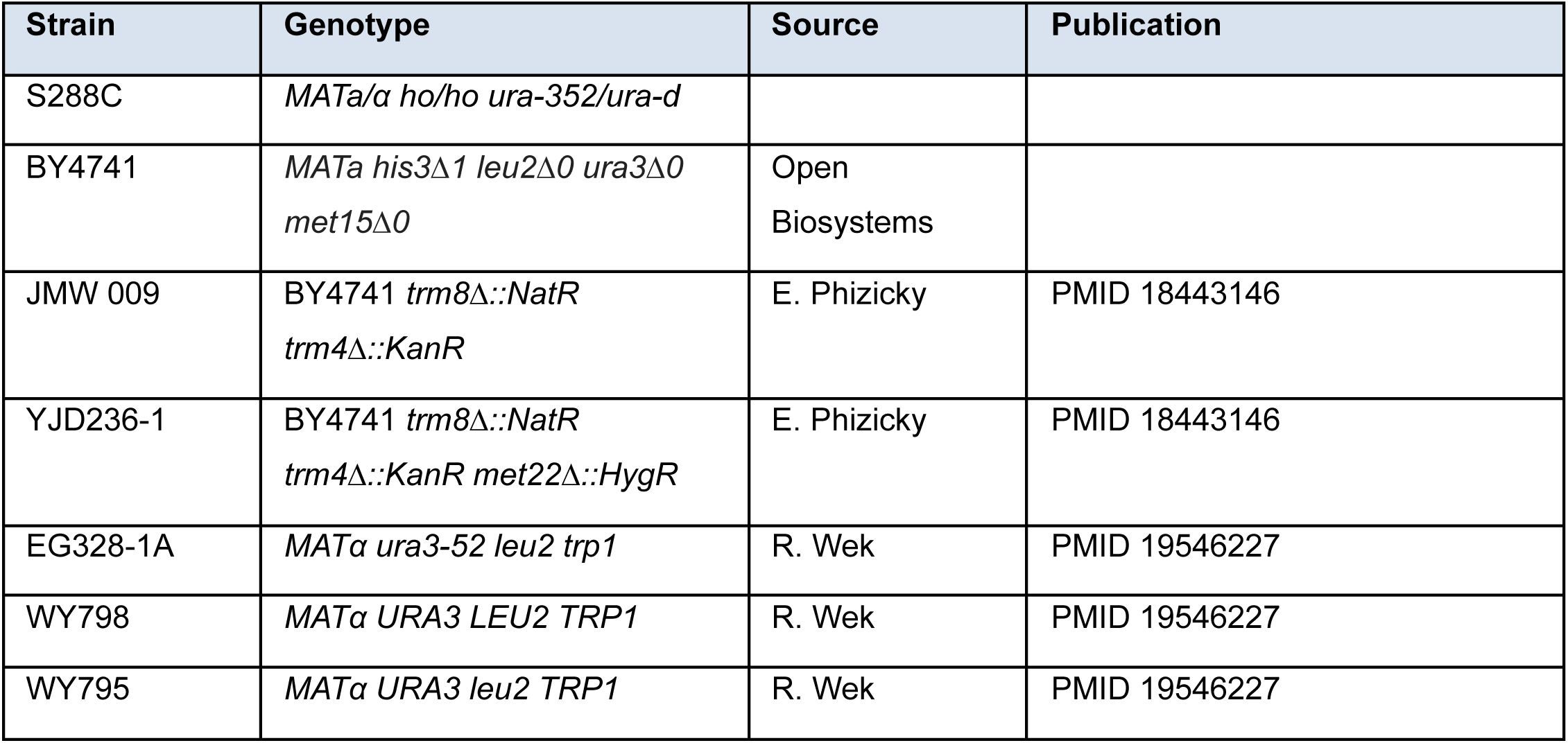
Yeast strains.

For validation of chemical charging northerns (**Fig. 1C**), a single colony of WY798 was inoculated into synthetic complete media and incubated at 30°C overnight with rotation. This culture was diluted to 50 mL of synthetic complete media the next day and allowed to shake overnight at 30°C before dilution to an OD_660_ of 0.2 in 100 mL synthetic complete media. The culture was grown to log phase at 30°C, collected by centrifugation, and resuspended in 100 mL room temperature synthetic complete media. The culture was allowed to grow in the new media for 15 minutes, shaking at 30°C before it was pelleted, washed with water, flash frozen in liquid nitrogen, and stored at −80°C.

For the nutrient stress experiment (**Fig. 3B-D**), 3 colonies of WY795 were inoculated into synthetic complete media and incubated at 30°C overnight with rotation before dilution to an OD_660_ of 0.2 in 50 mL the same media. The cultures were grown to log phase shaking at 30°C before pellets were collected by centrifugation. Each pellet was washed by resuspension in a small volume of synthetic media lacking uracil, tryptophan, histidine, and leucine and split into two tubes. Cells were pelleted again and the wash media was removed. One pellet from each original culture was resuspended in 25 mL 30°C synthetic complete media and the other in 25 mL 30°C synthetic media lacking leucine. They were allowed to grow in the new media for 15 minutes, shaking at 30°C before they were pelleted, washed with water, flash frozen in liquid nitrogen, and stored at −80°C.

For the temperature stress experiment (**Fig. 3E-H**), 3 colonies of JMW 009 and 3 colonies of JMW 510 were inoculated into YEPD media and incubated at 30°C overnight with rotation before dilution to an OD_660_ of 0.2 in 60 mL YEPD media. The cultures were grown to log phase shaking at 30°C. At this point, 50 mL of culture was pelleted by centrifugation, washed with water, flash frozen in liquid nitrogen, and stored at −80°C. 40 mL of 40°C YP glucose media was added to the remaining 10 mL of culture and these cells were grown for 3 hours at 37°C. They were then pelleted, washed with water, flash frozen in liquid nitrogen, and stored at −80°C.

### Isolation and chemical ligation of aminoacylated tRNAs from budding yeast

#### A complete protocol for aa-tRNA-seq is available on Benchling

Yeast pellets were thawed on ice and resuspended in 400 µL of cold AES (10 mM NaOAc pH 4.5, 1 mM ETDA pH 8, 0.5% SDS). 400 µL of cold 25:24:1 acid phenol:chloroform:isoamyl alcohol was added. Samples were vortexed for 15 seconds and allowed to rest on ice for 20 minutes, vortexing every 5 minutes. They were then spun at 18000 g for 10 minutes at 40°C and the aqueous phase was moved to a new tube.

A 0.4X volume of Ampure XP beads (Fisher Scientific A63881) were added to 100 µL of aqueous phase. They were rotated for 2 minutes at RT and placed on a magnet until the beads had settled. The supernatant was moved to a new tube and quantified via nanodrop. Small RNAs were isolated from 100 µg of this supernatant using a Zymo Research RNA Clean and Concentrator kit (R1018) according to the manufacturer’s instructions. Dilution of the bead supernatant for the first step of the kit was done with 10 mM NaOAc pH 4.5, not with water. Small RNA was eluted in 30 µL of 10 mM NaOAc pH 4.5 and quantified via nanodrop. It was stored at −80°C.

Two 3′ DNA-RNA hybrid splint adapters were designed with different internal sequences, one to ligate to deacylated tRNAs (“uncharged 3′ adapter”) and the other to acylated tRNAs (“charged 3′ adapter”, see **Table 2**). A universal 5′ adapter was designed to pair with either of the 3′ adapters. Syntheses of these adapters were ordered from IDT and resuspended in water to a concentration of 2 mM. Adapters were run on a 1.5 mM 6% TBU (Tris, boric acid EDTA) V16 polyacrylamide gel with 10 nmol loaded per lane (10-well comb). Staining was not performed and UV shadowing was used to excise the adapters. Gel slices were cut into small pieces and rotated end over end in crush + soak buffer (300 mM NaOAc pH 5.5, 1 mM EDTA pH 8.0, 0.1% SDS) overnight at 4°C. Nucleic acids were precipitated with 100% ethanol and resuspended in water to a final concentration of 100 µM.

**Table 2.**
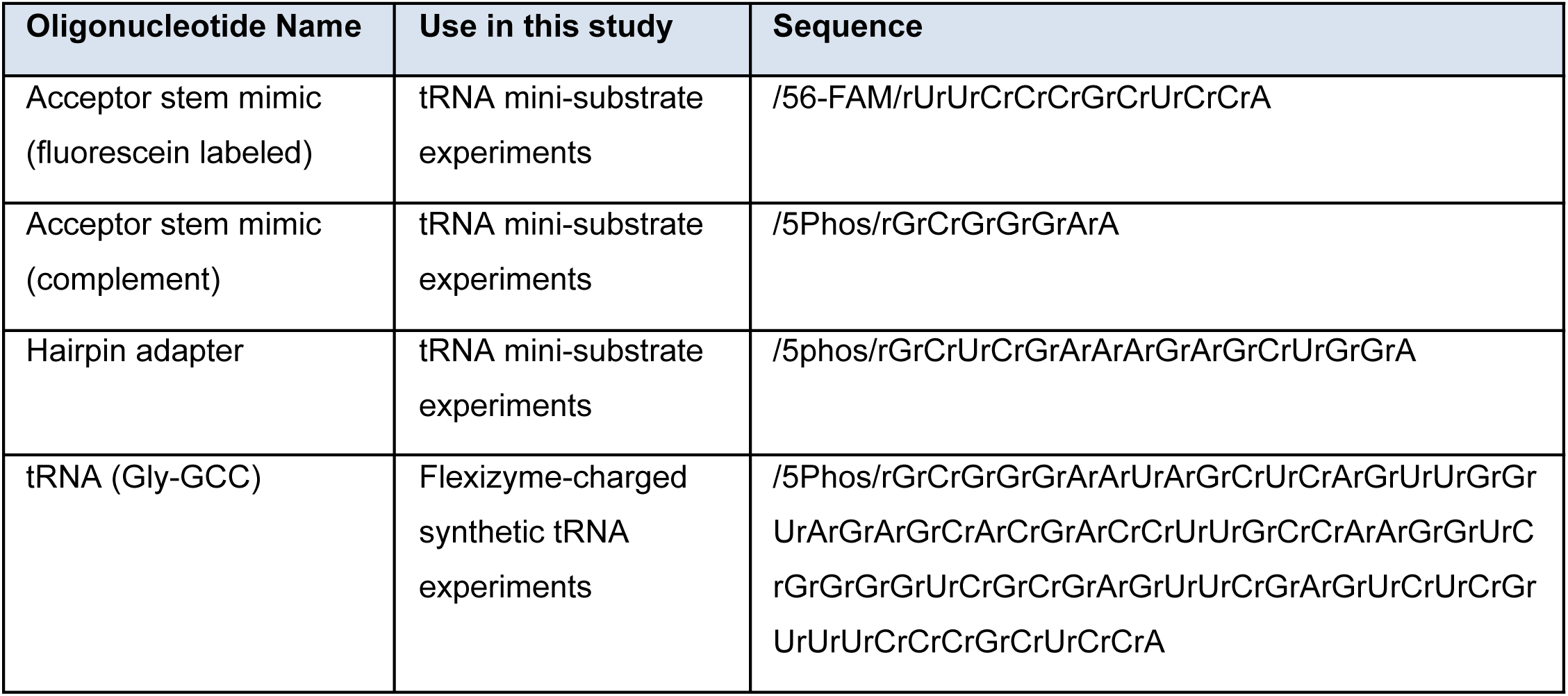
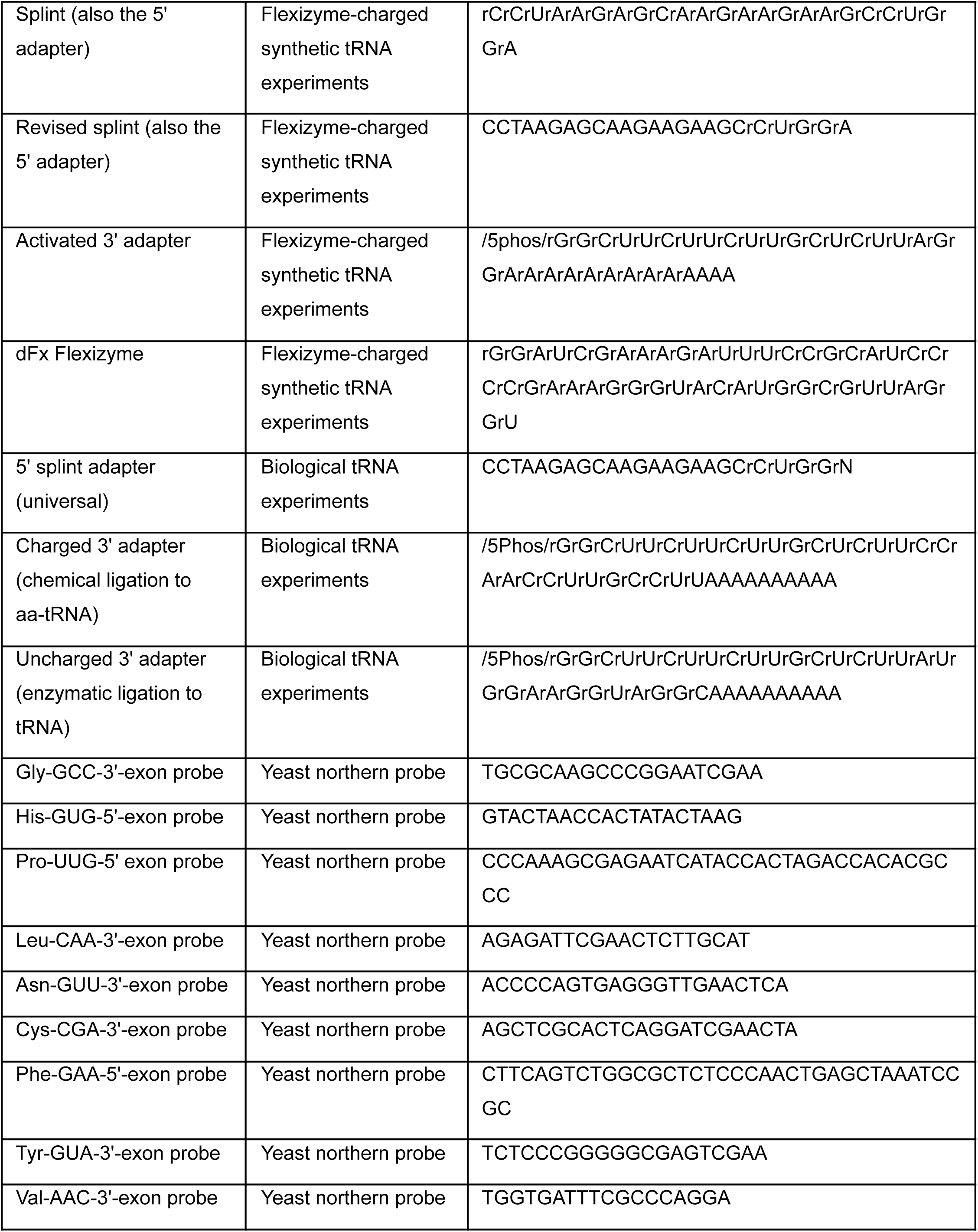
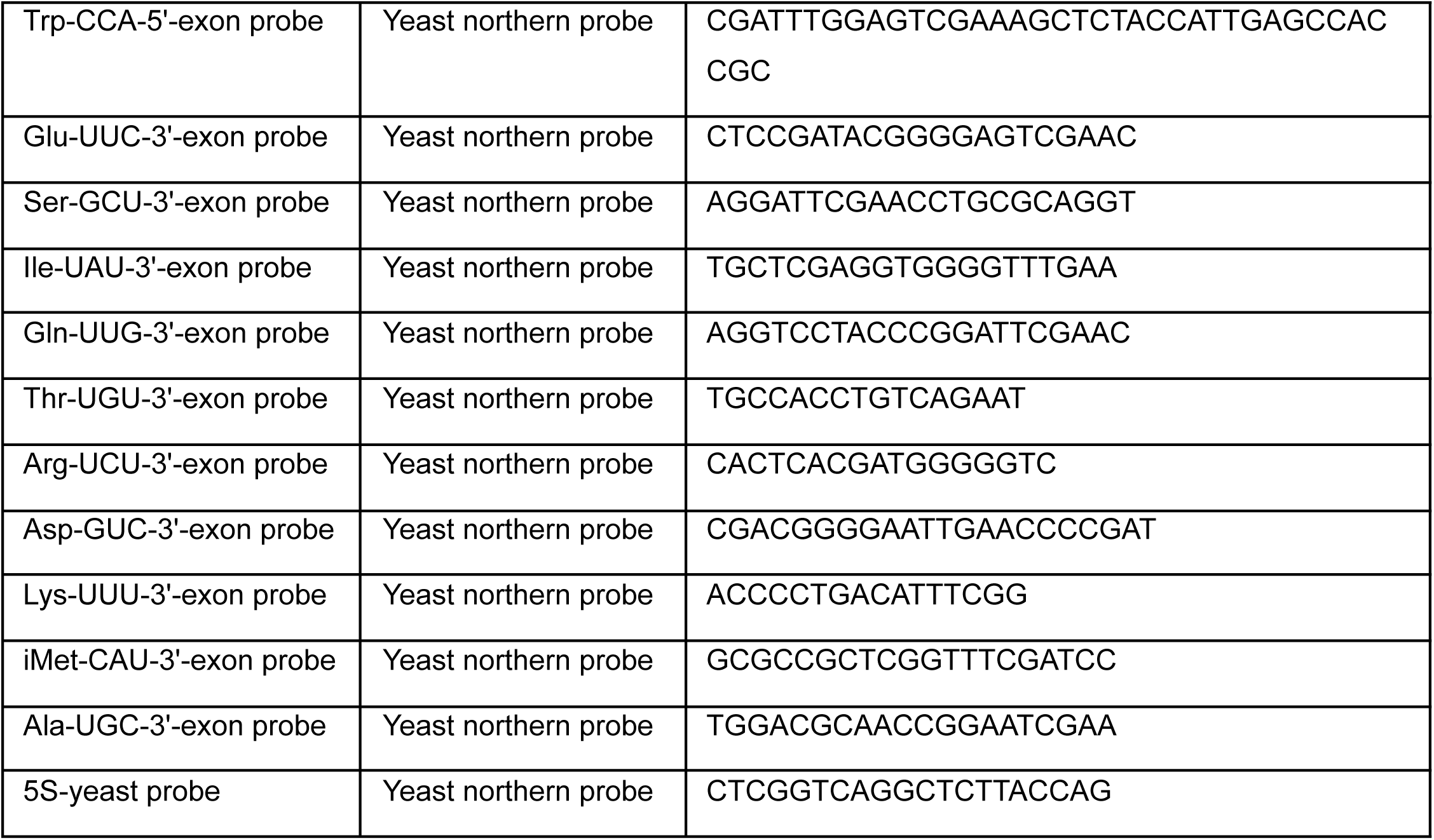
Oligonucleotide sequences.

Gel-purified charged 3′ adapter was incubated with a 500-fold molar excess of both imidazole and EDC (1-Ethyl-3-(3-dimethylaminopropyl)carbodiimide) for 2 hours at 25°C to imidazolate the 5′ end of the adapter. 30 µL of cold 99.5%+ acetone saturated with perchlorate and 1 mL of cold 99.5%+ acetone were added to precipitate the imidazolated adapter. The sample was incubated for 20 minutes on dry ice and then centrifuged at maximum speed for 10 minutes at 4°C. The supernatant was removed and the pellet was washed twice with 1 mL 1:1 acetone: diethyl ether. The pellet was dried in a speed vacuum and resuspended in 10 mM imidazole pH 7.0 to a final concentration of 200 µM.

Small RNA (15-50 pmol) was incubated in 100 mM MES pH 5.5, 2.5 mM MgCl_2_, a 5-fold molar excess of both imidazolated 3′ adapter and gel-purified 5′ adapter, and 50 mM HEI pH 6.5 for 30 minutes at 25°C establishing a phosphoramidate covalent linkage between the 3′ splint adapter and aminoacylated tRNAs. Ligated products were purified by crush and soak (0.3 M NaOAc pH 5.5, 1 mM EDTA pH 8.0, 0.1% SDS) at 4 °C, overnight) from a 10% TBU polyacrylamide gel, isolating the regions between 70 and 150 nts. The eluate was precipitated by addition of ethanol and GlycoBlue coprecipitant (Invitrogen) resuspended in a small volume of 10 mM NaOAc pH 4.5 and quantified via absorbance at 260 nm (Nanodrop).

##### Northern blotting

Transfer RNA charging was measured by acidic northern blot, resolving 75 ng of small RNA onto a gel (6% 19:1 acrylamide, 0.1 M sodium acetate, pH 4.5, 8 M urea) 42 cm in length which was run at 450 V for 22 hours in a cold room. For chemical-charging northern blots, chemically ligated tRNA (220 ng) was loaded onto 10% TBU polyacrylamide gels (7.5 cm, 6% 19:1 acrylamide, 1X TBE, 8 M urea) and electrophoresed in 1X TBE at room temperature at 250 V for 40 minutes. Acid-urea and TBU gels were transferred to charged nylon membranes (Hybond N+, GE) via electroblot transfer at 1 Amp for 1 hour for acid gels and 3 mA/cm^2^ based on the membrane area for 35 minutes for TBU gels. After transfer, membranes were UV-crosslinked at 254 nm using a 120 mJ dose and blocked in ULTRAhyb-Oligo (Thermo) before an incubation with ^32^P-labeled oligonucleotide probes in ULTRAhyb-Oligo overnight at 42°C (**Table 2**). Membranes were washed four times at 42°C (2X SSC, 0.1% SDS), wrapped in plastic, and exposed to a phosphor-imager screen before imaging on a Typhoon 9400 (GE Healthcare). Membranes were stripped with two 30 minute washes in 2% SDS at 80°C, prior to reblocking and incubation with labeled probe.

### Nanopore library generation for budding yeast tRNAs

Gel-purified tRNAs from chemical ligation were enzymatically ligated to capture deacylated tRNAs with 3′ splint adapters and attach 5′ adapters to all tRNA. tRNA from the first ligation (20 pmol) was incubated in a 20 µL reaction consisting of 10% PEG 8000, 1 µL of RNase inhibitor (Watchmaker Genomics), 9 pmol gel-purified uncharged 3′ splint adapter, 9 pmol gel-purified 5′ adapter, 1X T4 RNA ligase 2 buffer, and 2 µL of T4 RNA ligase 2 (homemade preparation, 0.74 mg/mL). This ligation was incubated at 25°C for 30 minutes.

Ligation products were purified by addition of a 1.8X volume of tRNA beads (BioDynami), mixing by pipetting, and incubation on ice for 4 minutes, followed by magnetic separation. The supernatant was discarded. Beads were washed with 180 µL 80% EtOH and air dried. Beads were resuspended in 13 µL of water, and the elution was moved to a new tube and quantified.

Splint-adapter-ligated tRNAs are next ligated to RT adapters (RTA) (provided in the RNA004 ONT kit): 12.5 µL of sample was incubated with 1.5 µL RTA, 0.5 µL RNase inhibitor (Watchmaker Genomics), 4 µL T4 DNA ligase buffer, and 1.5 µL T4 DNA ligase (Watchmaker Genomics) for 30 minutes at 25°C, and cleaned up at RT using the tRNA beads as above, using a 1.35X volume of beads, and elution in 26 µL water. Each sample was quantified with the Quant-iT Qubit dsDNA HS kit and the library size distribution was confirmed by Agilent TapeStation (HS DNA 1000).

Finally, ligation products were ligated to ONT’s RNA ligation adapter (RLA) on the same day that sequencing was conducted. 50-400 fmol of sample in 23 µL was incubated with 6 µL RLA (ONT RNA004), 8 µL T4 DNA ligase buffer, and 3 µL T4 DNA ligase (Watchmaker Genomics) for 30 minutes at 25°C. These final ligations were cleaned up using a 1.8X volume of Ampure XP SPRI beads (Beckman Coulter), and washed with WSB wash buffer (ONT) following the protocol for ONT SQK-RNA004.

### Sequencing run conditions for budding yeast tRNAs

Libraries were loaded onto “RNA” flow cells on a PromethION P2 Solo instrument connected to a A5000 GPU workstation or a PromethION P2integrated instrument, using MinKNOW software version 24.06.10. We found the throughput of aa-tRNA-seq to be comparable or superior to previous nanopore tRNA sequencing approaches (*32*, *76*), collecting a median >8 million reads for biological tRNA sequencing libraries, and median ∼250 thousand reads for synthetic tRNA sequencing libraries.

### Base-calling and alignment

Libraries were basecalled with Dorado v0.7.2 (Oxford Nanopore Technologies, https://github.com/nanoporetech/dorado) using the “super high accuracy” (rna004_130bps_sup) v5.0.0 model and –emit-moves parameter. Basecalled bams were converted to fastq format using samtools v. 1.21 (*77*) with -T “*” flag to retain move tables, and then aligned to using BWA-MEM version 0.7.16-r1181 (*78*) with the parameters `bwa mem -C -W 13 -k 6 -x ont2d`, enabling transfer of move tables to aligned bams.

To evaluate the performance of an alignment-based classification approach for identifying charged vs. uncharged tRNA reads, aligned bams were further filtered to contain reads mapped to full length tRNAs. tRNA reference files were constructed by appending CCA sequences to each mature tRNA sequence, along with the unique 3′ and universal 5′ adapter sequences, and primary alignments for each read assessed. A Snakemake (*79*) analysis pipeline will be available upon manuscript acceptance at https://github.com/rnabioco/aa-tRNA-seq-pipeline.

### Remora model training and validation

To train a machine learning model distinguishing charged and uncharged tRNAs we used the Remora software (v3.2, https://github.com/nanoporetech/remora) and training procedure for modified nucleosides. First, we prepared fully charged yeast tRNA libraries (treated as modified base in the training) and deacetylated libraries (treated as modified base control). To make our model universal for all tRNAs we defined a 6-nt modification kmer spanning the universal CCA 3′ end of tRNA and the first three nucleotides of the 3′ adapter (CCA<u>G</u>GC), where the underlined G was defined as the modification site. We extract chunks from both libraries using `remora dataset prepare` with the default remora 9mer table for rescaling and following parameters: `--refine-rough-rescale –reverse-signal –motif CCAGGC` 3. We prepared training configuration files with `remora dataset make_config using –dataset-weights 1 1`. Finally we trained the model using the parameters: `--model ConvLSTM_w_ref.py –chunk-context 200 200 --num-test-chunks 20000`. After internal remora validation on 20000 chunks, this model was validated on independent libraries using reference-anchored remora inference (`remora infer from_pod5_and_bam –reference-anchored`). We used ML tags from inference output bams to calculate model performance, with the following assumption: ML < 200 = uncharged tRNA, ML ≥ 200 = charged tRNA.

Pairwise machine learning classifiers to distinguish individual amino acids (binary recognition between pairs of amino acids totalling 380 models) were trained using Remora v. 3.2 using a procedure similar to that described above, using Flexizyme-charged synthetic tRNA reads for the training. Synthetic tRNA was aligned to a reference sequence with one nucleotide (“T”) inserted between the CCA sequence at the 3′ terminus and the start of the 3′ adapter sequence. Pairwise models were trained with one amino acid treated as modified base and the second one as modified base control on the CCAT motif, with the inserted T identified as the modification position for Remora training. 10000 chunks were used for internal Remora validation, except for pairwise comparisons with alanine, where 9000 chunks were used due to lower library depth for the alanine-charged library. F1 scores were calculated for all models using the equation: `2TP/(2TP+FP+FN)`.

### Nanopore signal analysis

Signal metrics (dwell time and trimmed means of ionic current) at reference-anchored positions were extracted from POD5 files using the Remora API and stored in TSV files for analysis and plotting in R. Example scripts for signal extraction and plotting will be available at https://github.com/rnabioco/aa-tRNA-seq.

## Acknowledgements

This work was supported by the National Institutes of Health (R35 GM119550) and National Science Foundation (Award #2330283). J.W.S. is an Investigator of the HHMI. We thank Filip Bošković for inspiring discussions during the project planning phase, Ron Wek and Eric Phizicky for yeast strains, Marcus Stoiber for advice and assistance with Remora model training, and members of the Hesselberth and Szostak labs for constructive feedback.

## Data availability

Complete sequencing data (including POD5 and FASTQ files) and classification models will be available at the NCBI Gene Expression Omnibus and Zenodo upon manuscript acceptance. Details on the associated BioProject will be included on both the GitHub repository (https://github.com/rnabioco/aa-tRNA-seq) and this manuscript upon acceptance.

---

**Figure S1.**
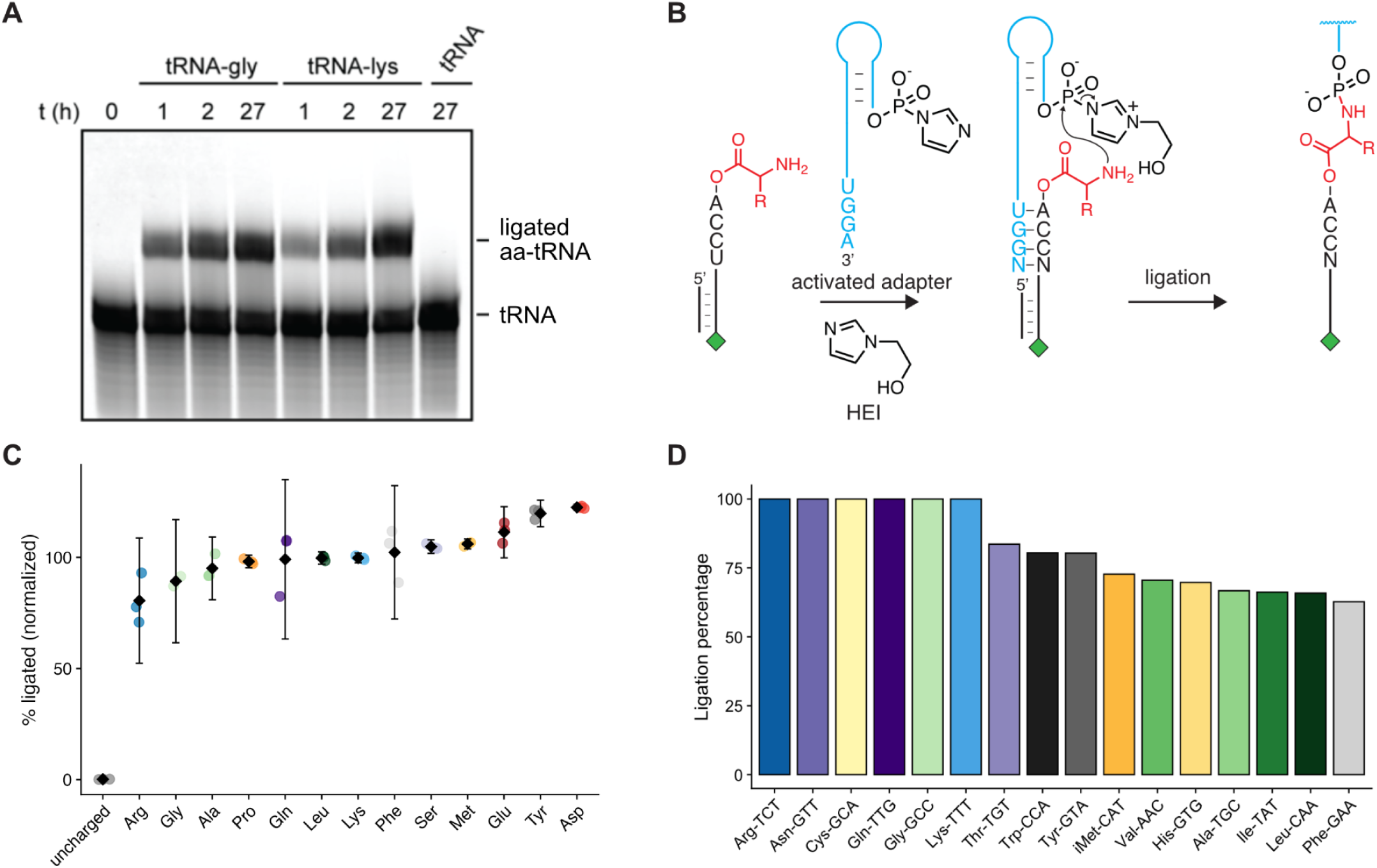
(**A**) Chemical ligation of Flexizyme-charged synthetic tRNA in the absence of catalyst for 0, 1, 2, or 27 hours. The analytical 16 % denaturing gel was stained with SYBR gold to visualize the RNA. (**B**) Schematic of an nt tRNA minisubstrate bearing a fluorophore (FAM, green diamond) undergoing chemical ligation to 5′-phosphorimidazolide activated hairpin adapter in the presence of the catalyst 1-(2-hydroxyethyl)imidazole (HEI). (**C**) Normalized levels of ligation product for fluorescently labeled tRNA minisubstrate in (**B**) aminoacylated with the indicated amino acids using Flexizyme and reacted with the activated hairpin adapter in the presence of HEI for 90 minutes. (**D**) Densitometry-based quantifications of the percent of aminoacylated tRNA shifted upon chemical ligation for the budding yeast tRNAs visualized on the acidic northern in **Fig. 1B**, after stripping and reprobing for the indicated isodecoders.

**Figure S2.**
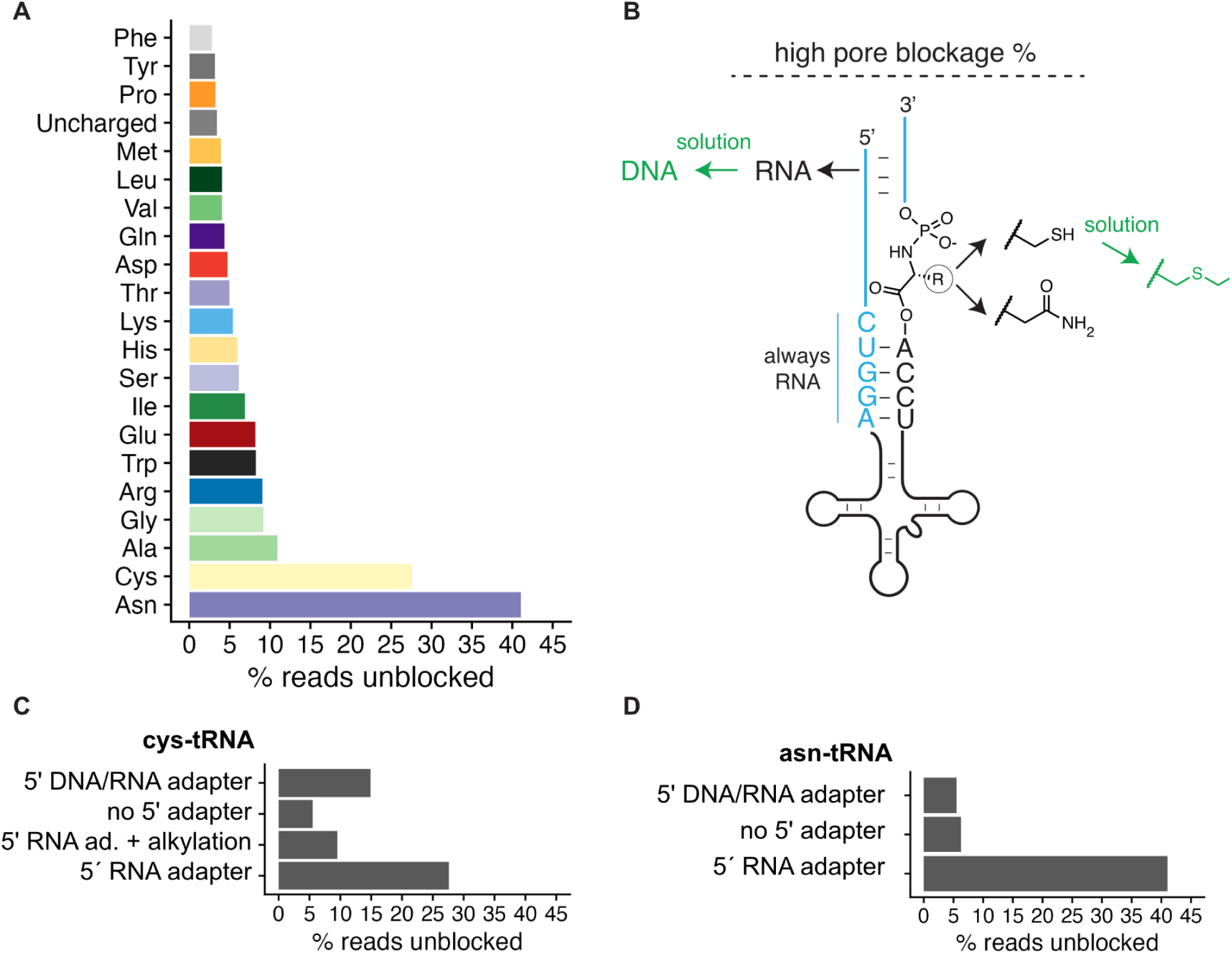
(**A**) Percentage of reads with "unblock" end status in nanopore sequencing libraries from synthetic tRNA chemically ligated to an imidazolated 3′ RNA adapter and enzymatically ligated to a 5′ RNA adapter. (**B**) Schematic depicting experimental strategies tested for their effects on Asn- and Cys-tRNA pore blocking rates, including incorporation of a DNA/RNA hybrid 5′ adapter, alkylation of cys-tRNA, or hydrolysis of Flexizyme-charged asn-tRNA. (**C**) Percent of Cys-tRNA reads with "unblock" status using each of the relevant strategies above, as well as omission of 5′ adapter. (**D**) Equivalent percentage of "unblocked" Asn-tRNA reads for all three approaches tested.

**Figure S3.**
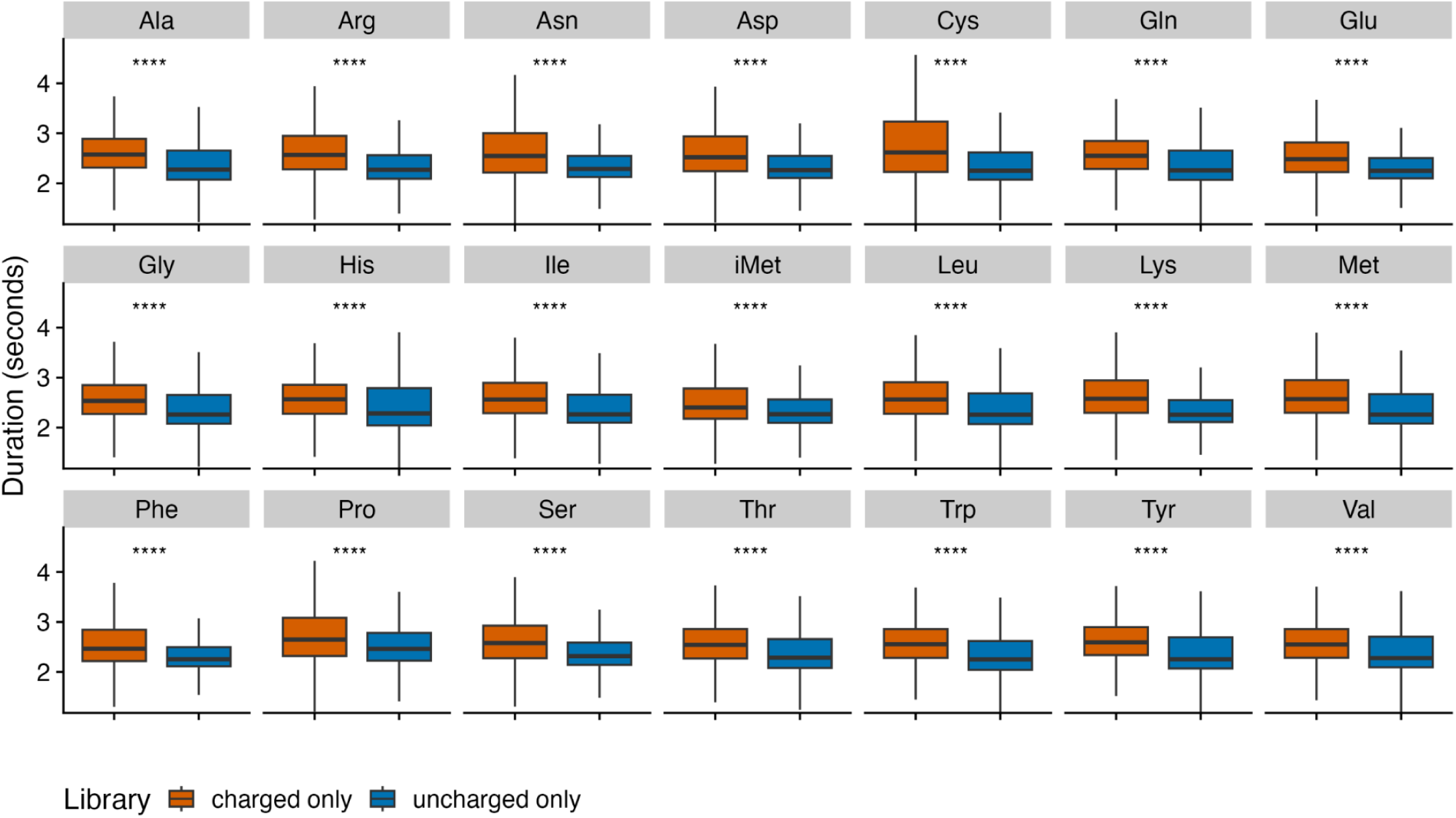
tRNA translocation time by isodecoder in budding yeast tRNA sequencing libraries prepared using only chemical ligation to imidazole-charged adaptors ("charged only", orange), or libraries where tRNA was first chemically deacylated followed by enzymatic ligation with T4 RNL2 ("uncharged only", blue). The y-axis displays the translocation duration for the entire read in seconds. Statistical significance between the distributions in each library was assessed using the Wilcoxon test, with significance levels indicated by asterisks above each comparison (****p < 0.0001).

**Figure S4.**
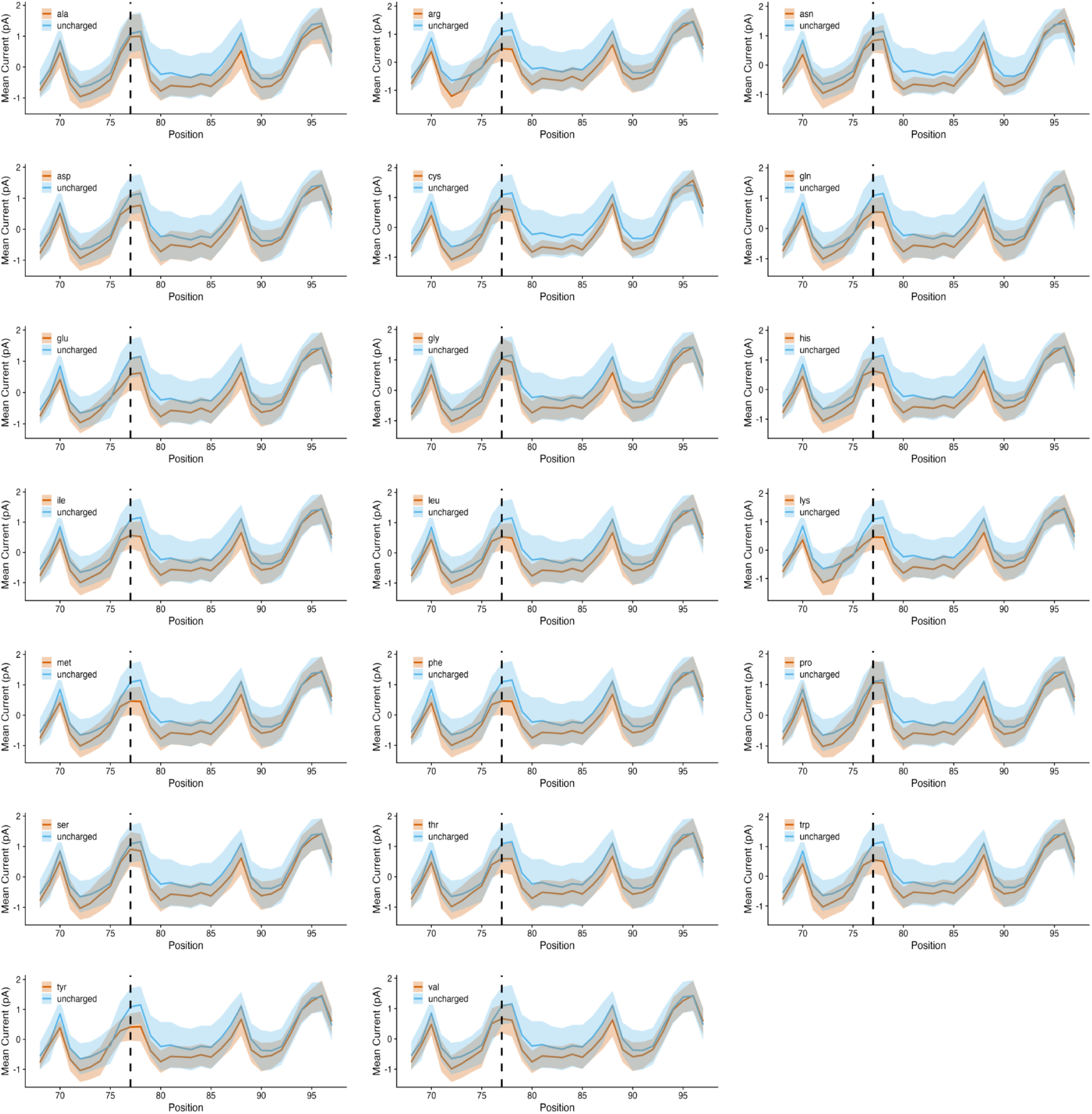
Normalized signal intensity (current in picoamps) for synthetic tRNA charged with 20 naturally-occurring amino acids using Flexizyme. Each panel plots the mean current in picoamps for an uncharged tRNA substrate in the central blue line, with the shaded blue area representing the standard deviation around the mean. Each aminoacylated comparison is plotted analogously in orange. The x-axis spans the same window of interest as in **Fig. 2**, containing six nucleotides at the 3′ terminus of the tRNA, the CCA tail, the aminoacylated position (dashed line, included as an extra nucleotide inserted at position 77 in the reference) and the entirety of the 3′ adapter sequence from nt 78 onward.

**Figure S5.**
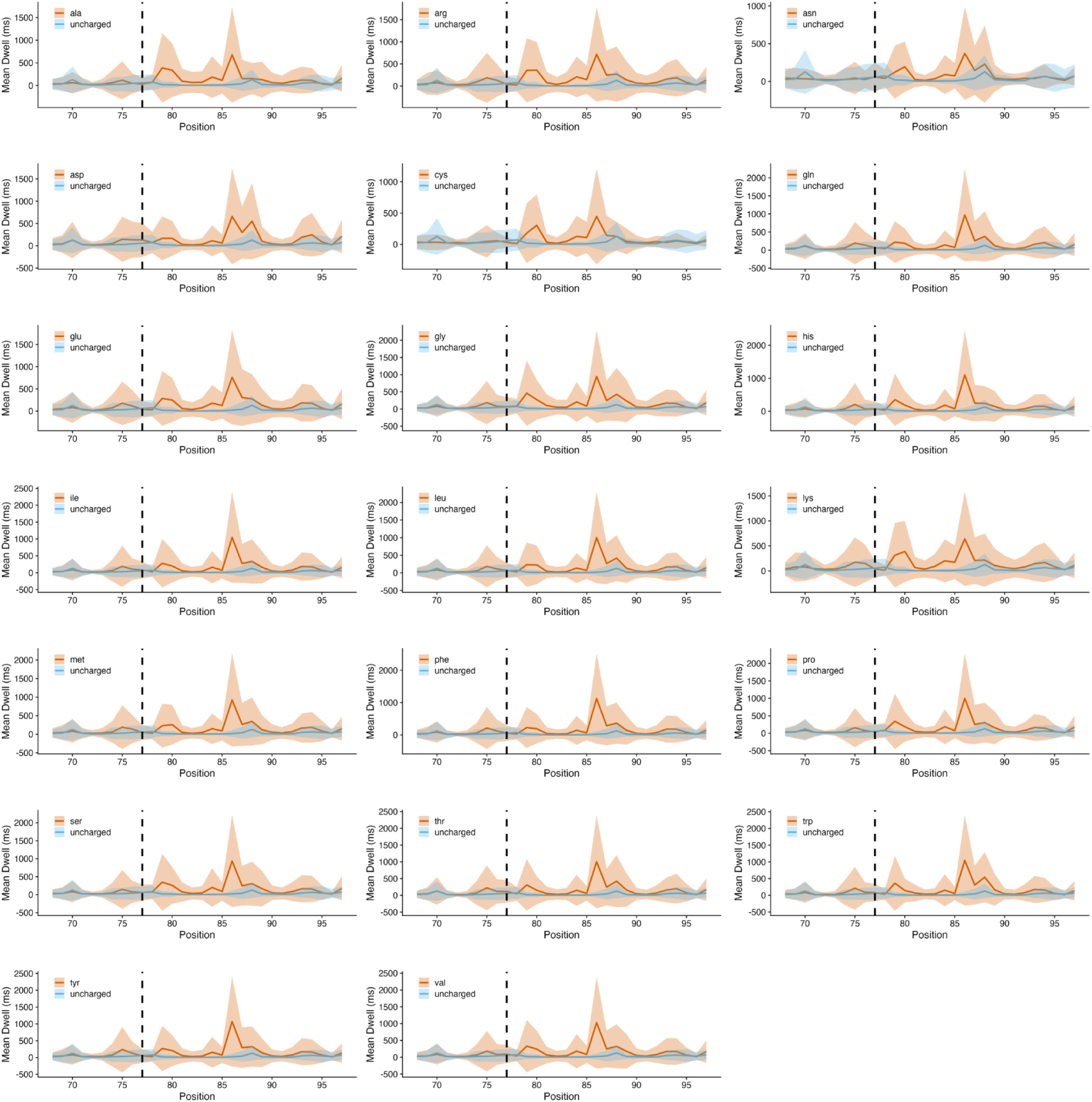
Mean dwell time for synthetic tRNA charged with 20 naturally-occurring amino acids using the Flexizyme. Each panel plots the mean dwell in milliseconds at each nucleotide in an uncharged tRNA substrate in the central blue line, with the shaded blue area representing the standard deviation around the mean. Each aminoacylated comparison is plotted analogously in orange. The x-axis spans the same window of interest as in **Fig. 2**, containing six nucleotides at the 3′ terminus of the tRNA, the CCA tail, the aminoacylated position (dashed line, included as an extra nucleotide inserted at position 77 in the reference) and the entirety of the 3′ adapter sequence from nt 78 onward.

